# Large-scale color biases in the retinotopic functional architecture are region specific and shared across human brains

**DOI:** 10.1101/2020.05.31.126532

**Authors:** Michael M. Bannert, Andreas Bartels

## Abstract

Despite the functional specialization in visual cortex, there is growing evidence that the processing of chromatic and spatial visual features is intertwined. While past studies focused on visual field biases in retina and behaviour, large-scale dependencies between coding of color and retinotopic space are largely unexplored in the cortex. Here we asked whether spatial color biases are shared across different human observers, and whether they are idiosyncratic for distinct areas. We tested this by predicting the color a person was seeing using a linear classifier that has never been trained on chromatic responses from that same brain, solely by taking into account: (1) the chromatic responses in other individuals’ brains and (2) commonalities between the spatial coding in brains used for training and the test brain. We were able to predict the color (and luminance) of stimuli seen by an observer based on other subjects’ activity patterns in areas V1-V3, hV4 and LO1. In addition, we found that different colors elicited systematic, large-scale retinotopic biases that were idiosyncratic for distinct areas and common across brains. The area-specific spatial color codes and their conservation across individuals suggest functional or evolutionary organization pressures that remain to be elucidated.

**Significance statement:** Does a color elicit comparable neural activity in two different observers? We addressed these questions, which involved the central test to predict what color someone is seeing based on their brain activity if it is only known how other brains respond to color. We estimated the commonalities across brains in the way they respond to achromatic, spatial stimulation, which allowed us to retinotopically align different brain responses to each other in a common response space. In this space derived without any color responses we could decode across brains what color an observer was seeing and found that spatial color biases differed between areas. Our results demonstrate systematic dependencies between chromatic and visual field representations that are area-specific and preserved across observers.

## Introduction

It is known that the primate visual cortex contains multiple representations of the visual field in different functional areas (Felleman & Van Essen, 1991; Sereno et al., 1995; Wandell et al., 2007). This multiplicity has been explained in terms of differences between areas in what cognitive and behavioral purpose their computations fulfill (Goodale & Milner, 1992; Mishkin & Ungerleider, 1982), what perceptual processes they reflect (Gauthier et al., 2000), what content or object properties they encode (Kanwisher et al., 1997; Malach et al., 1995; Tanaka, 1993; Zeki et al., 1991).

While differences between brain areas are widely accepted, within-area variability that can be explained by the underlying cortical retinotopy received less attention (Freeman et al., 2011, 2013; Roth et al., 2018; Sasaki et al., 2006). Since these few studies focused on orientation tuning only, even less is known about color.

Traditionally, human color vision has been thought to be mediated by functionally segregated mechanisms already at earliest processing stages (De Valois & Jacobs, 1968; Derrington et al., 1984; Field & Chichilnisky, 2007; Schiller et al., 1990; Zeki, 1973). Independent processing could explain dissociations in LGN and visual cortex, with a specialization for color processing in the cytochrome-oxidase rich “blob” and “thin stripe” regions in areas V1 and V2, respectively (Denison et al., 2014; Livingstone & Hubel, 1984, 1988; Lu & Roe, 2008; Nasr et al., 2016; Schiller et al., 1990; Ts’o & Gilbert, 1988), and in V4 (Bartels & Zeki, 2000; Beauchamp et al., 1999; Lueck et al., 1989; Zeki, 1978, 1980; Zeki et al., 1991), especially its “globs” (Conway et al., 2007; Conway & Tsao, 2009).

There are a few neuroimaging experiments investigating how color processing in the cortex differs across the visual field, but unfortunately they have been limited to V1 and investigated the radial field coordinate only (Mullen et al., 2007; Vanni et al., 2006). This may surprise considering the inhomogeneities in retinal cone distributions (Roorda & Williams, 1999) and psychophysical evidence for differences in chromatic sensitivity as a function of visual space (Danilova & Mollon, 2009; Horiguchi et al., 2013; Levine & McAnany, 2005; Mullen, 1991; Mullen & Kingdom, 2002). Computational analysis furthermore suggested that the difference in color statistics between objects and surrounding background contributes to color tuning in object recognition regions (Rosenthal et al., 2018). It thus makes sense that objects, which are typically foveated, might benefit from different chromatic processing than the peripheral background. Finally, simulations showed that, to explain visual performance differences across the visual field, retinal factors are insufficient, suggesting cortical contributions are plausible (Kupers et al., 2019).

Our research aim was to examine functional differences in color processing as a function of visual space in retinotopic areas. Specifically, we were interested to learn how consistent such dependencies would be across different brains.

We addressed these questions by aligning subject-specific fMRI response spaces to each other using shared response modeling. In contrast to previous similar cross-subject alignment approaches (Guntupalli et al., 2016; Haxby et al., 2011), we used brain responses evoked by the spatially changing position of slowly moving achromatic checkerboard stimuli used for retinotopic mapping. Individual brain activity was hence sampled in response to a highly restricted subset of stimulus inputs comprised of purely spatial variability in the absence of chromatic stimulation. This ensured that the stimulus properties that were used for the estimation of shared neural responses were based solely on spatial information. If neural co-representations of space and color are indeed shared across observers, it should be possible to predict from an observer’s pattern of brain activity what color the person was seeing using a classifier that was trained solely on other observers’ brain responses to color, i.e., to classify color across the brains of different participants. Our findings showed this to be true, and further suggest that the observed between-subject decoding is mediated by common spatio-chromatic representations that are at least partially embodied by large-scale retinotopic biases shared between brains yet idiosyncratic for distinct regions in the visual processing hierarchy.

## Materials and Methods

### Participants

We analyzed fMRI data from N = 15 (2 male, 13 female) participants aged between 22 and 35 years (mean: 25.5) who took part in a previously published fMRI study about color vision (Bannert & Bartels, 2018). We only selected participants from that study for which the cortical retinotopic representations of the visual field were measured along both the polar and the eccentricity axis of the visual field (see below). All participants had normal or corrected-to-normal visual acuity and were tested for normal color vision using Ishihara color plates (Ishihara, 2011). Each participant gave written informed consent before the first study session. The experiment was approved by the local ethics committee of the Tübingen University Hospital.

### Experimental setup

All stimuli were shown to the volunteers while they were lying in the scanner using a projector (NEC PE401H) that was gamma-calibrated with a Photo Research PR-670 spectroradiometer running the Psychtoolbox-3 display calibration script. Experimental stimuli were presented at a resolution of 800×600 pixels and a refresh rate of 60 Hz. The display on the projection screen had a size of 21.8° by 16.2° of visual angle. Stimuli were presented using Psychtoolbox-3 (Kleiner et al., 2007) and MATLAB running on Windows XP.

### Color Stimuli

Brain responses obtained from a retinotopic mapping session and a color vision experiment were used for the present analysis. Stimuli of both sessions have been described in detail before (Bannert & Bartels, 2018), and are again briefly described below.

In the color vision experiment, we showed stimuli with each belonging to one of three different color categories: red (mean chromaticities x = 0.39, y = 0.35), green (mean chromaticities x = 0.34, y = 0.41), yellow (mean chromaticities x = 0.41, y = 0.43). Each color category was shown at a high or low luminance, resulting in a total of six trial types. Each stimulus consisted of concentric rings presented against a mid-level gray background (154 cd/m^2^). Stimulus intensities were psychophysically matched (see below) for the high and low luminance conditions, respectively, yielding mean luminance values (SD in brackets) of 241.7 (19.8) cd/m^2^ and 199.8 (9.0) cd/m^2^ for red, 242.4 (11.4) cd/m^2^ and 190.0 (8.8) cd/m^2^ for green, 230,4 (11.6) cd/m^2^ and 179.6 (9.8) cd/m^2^ for yellow. Transparency of the stimulus color was sinusoidally modulated as a function of radial distance from the center (thereby conferring the appearance of multiple concentric rings). The radius of the largest ring was 8.61° of visual angle and the cycle size of the modulation was 2.16° of visual angle with its phase changing continuously at a speed of 2.47°/s in an outward direction. To ensure that the three colors appeared isoluminant, we used the minimum flicker method (Kaiser, 1991) inside the scanner prior to data acquisition: a color rectangle of size 3.28° by 2.46° of visual angle in vertical and horizontal directions from a given color category was shown foveally to the participant while continuously replacing it every second frame with an achromatic reference stimulus of a given luminance. By adjusting the luminance of the color patch by button press in steps of 11 – 12 cd/m^2^, the participant chose the stimulus intensity that minimized the amount of perceived flicker. Since the three color conditions were presented in either low or high luminance (151.3 cd/m^2^ or 184.9 cd/m^2^), each participant individually performed adjustments with reference stimuli at two different luminance levels. Our color experiment thus had a 3-by-2 factorial design.

### Experimental design and task

The observers had to foveate the fixation dot in the middle of the screen while paying attention to the expanding color rings. Each stimulus was presented for 8.5 s followed by an inter-trial-interval (ITI) of 1.5 s. There were 36 trials per imaging run for a total of 216 trials presented across the six runs. The trial sequence was pseudo-randomized across runs to ensure that each of the 6 conditions was preceded equally often by every condition (Brooks, 2012). The last trial in every run was repeated at the beginning of the subsequent run to ensure that the trial sequence remained counterbalanced across all runs. The first trial of each run was excluded from the analysis.

The task was to indicate by button press the occurrence of a brief (0.3 s) luminance increment or decrement of the color stimulus. Specifically, when presenting a high luminance color stimulus, the target was the low luminance stimulus and vice versa. Observers were instructed to respond as quickly and accurately as possible and received visual feedback about mean reaction time (RT) and errors at the end of each imaging run.

### Retinotopic mapping details

In the retinotopic mapping experiment each participant underwent four runs of polar angle and two runs of eccentricity mapping following standard protocol (Sereno et al., 1995; Wandell & Winawer, 2011). In polar runs observers viewed an achromatic contrast-reversing checkerboard stimulus through a wedge-shaped aperture in a mid-level gray layer occluding the checkerboard. The wedge had an angle of 45 degrees and extended to the edge of the display. The aperture slowly rotated in a clockwise direction in half of the runs or counter-clockwise direction in the remaining half. In the two eccentricity runs the checkerboard stimulus was viewed through an annulus that cyclically expanded in one run whereas it contracted in the other. The check sizes in the polar and eccentricity runs as well as the annulus width increased logarithmically with radial distance to accommodate cortical magnification. Participants were instructed to keep their eyes on the fixation cross in the middle of the screen while paying attention to the wedge or annulus apertures, respectively, and to press a button each time a red dot briefly appeared randomly anywhere in the visible checkerboard.

Per run, most participants saw 10 cycles of the rotating wedge and expanding or contracting annulus with each cycle lasting 55.68 s. For subjects 2, 8, and 14, the stimulus period was 50.46 s and 12 cycles were presented. Subject 1 saw only 9 cycles. To be able to use shared response modeling (see below), we made sure that the stimulus input was the same for each time point across participants by including only fMRI measurements for the first 9 cycles and temporally resampling data from subjects, 2, 8, and 14 to obtain 576 fMRI volumes for each participant. However, we used each participant’s original dataset to define retinotopic regions-of-interest (ROIs) and determine polar coordinates for each voxel.

### fMRI scan details

Imaging was carried out on a Siemens Prisma scanner at 3 T magnetic field strength using a 64 channel head coil. The 56 slices were positioned, without gaps between them, approximately parallel to the AC-PC line for whole brain coverage. We employed a multi-band factor 2 and GRAPPA factor 2 to obtain a four-fold accelerated parallel imaging sequence. T2*-weighted functional images were recorded at a repetition time (TR) of 0.87 s and an in-plane matrix resolution of 96×96. Slice thickness and voxel size were 2 mm isotropic. Echo time (TE) was 30 ms and flip angle was 57°. Anatomical images were recorded for each participant at a voxel size of 1 mm isotropic using a T1 weighted MP-RAGE ADNI sequence. To correct for magnetic field inhomogeneities, we measured gradient field maps which were included in the motion correction step of the fMRI preprocessing pipeline.

### fMRI data preprocessing

Data from the main experiment were preprocessed using SPM8 (https://www.fil.ion.ucl.ac.uk/spm) running on MATLAB 2014b (The Mathworks, Inc., Natick, MA, USA). We excluded the first 11 images of each run to allow the magnetic field to reach equilibrium. To correct for head motion, each participant’s functional images were realigned to the first recorded image. The gradient field maps were used to unwarp the image sequence in order to take into account magnetic field distortions. We corrected for differences in slice acquisition times by shifting the phase of each frequency in the signal’s Fourier representation to the middle of the volume. Every participant’s image sequence was then co-registered to their respective anatomical scan. Anatomical scans were spatially normalized to MNI space using SPM’s segmentation-based method and the ensuing transformations were also applied to normalize functional images to MNI space.

The data from the retinotopic mapping experiment were motion-corrected, co-registered, and slice-time-corrected with SPM8 in the same way as the data from the main experiment. We used FreeSurfer’s recon-all pipeline (https://surfer.nmr.mgh.harvard.edu) to reconstruct each participant’s cortical surface from their individual anatomical scan (Fischl, 2012). The functional images were then spatially smoothed on the cortical surface with a 4 mm Gaussian kernel.

### Retinotopic ROI definition

We used standard retinotopic procedures to functionally identify visual areas in each participant separately (Sereno et al., 1995; Wandell & Winawer, 2011). After applying Fourier-transformation to the time series of each vertex, we plotted the corresponding phase of each vertex at the frequency of the polar angle mapping stimulus onto the inflated cortex. The reversals in the angle map indicate the boundaries between visual areas. In this way, we delineated boundaries for areas V1, V2, V3, hV4, VO1, LO1, and LO2.

### Pattern estimation and preparation

To extract patterns of brain activity we fit each voxel time series with a general linear model (GLM) in SPM8. The design matrix contained one boxcar regressor for each of the 216 trials (36 trials per run, six runs in total), a regressor each for the first trials in every run, which were excluded from the analysis, a constant intercept for each run, and the six parameters from the motion correction step to model out motion-induced artifacts in the voxel time series. To account for the hemodynamic lag of the BOLD response, we shifted each boxcar 5 s forward in time. The beta weights were obtained using restricted maximum likelihood estimation and were then used to form vectors of brain responses to their respective color stimulus. Finally, before a sequence of vectors were entered into pattern classification analyses, each voxel time series within this sequence was – for each run separately – temporally detrended by removing the fit of a second-order polynomial (effectively high-pass filtering the data), followed by scaling to zero mean and unit variance

### Pattern classification

Pattern classification was performed in Python 3 using scikit-learn 0.19.1 (Pedregosa et al., 2012). We used Linear Discriminant Analysis (LDA) to train classifiers based on a shrinkage estimate of the covariance matrix (Ledoit & Wolf, 2004). In a first step, we tested how well color (and luminance) information could be decoded from brain signals when the training and test data came from the same participant. In the second step, we examined how well color (or luminance) patterns generalized across participants. We refer to the analyses in the first and second steps as within-subject classification (WSC) and between-subject classification (BSC), respectively (Haxby et al., 2011).

**In WSC analyses,** we trained LDA classifiers to predict from vectors of neural responses from which of the three color categories (or the two luminance conditions in the luminance classification) they came. We obtained unbiased estimates of classification performance by cross-validating them following a leave-one-run-out cross-validation procedure. There were 6 runs with 36 response vectors each. The classifier was thus trained on 180 vectors (5 times 36) and tested on 36 vectors in every iteration. Each vector element corresponded to the response estimate of a single voxel within a given ROI. Individual results were averaged across all participants.

**In BSC analyses,** however, we were interested to learn how well color (or luminance) responses generalized across participants. Rather than cross-validating classifiers across runs (within each participant separately) and then averaging results over participants we now cross-validated across participants and obtained a single value for the whole group. Another difference was that vector elements used for classification came from the *k*-dimensional common space estimated through shared response modeling (see below) (instead of voxels used in WSC) for each ROI. With 216 response vectors for each of our 15 participants, this leave-one-participant-out cross-validation approach resulted in classifiers being trained on 3024 (14 times 216) response vectors and tested on the 216 response vectors of the left-out participant in every iteration. The *k* vector elements corresponded to the common space dimensions, i.e. weighted sums of the voxel estimates from individual observers’ ROIs. The value of *k* was determined via grid search on the training set for each ROI (see **Shared response modeling**). Since shared response modeling implicitly already performs dimensionality reduction, we did not use any additional feature selection such as recursive feature elimination.

Since color decoding was a three-way classification whereas luminance decoding was a two-way classification, classification accuracies were transformed to z-values using the normal approximation to the binomial distribution:

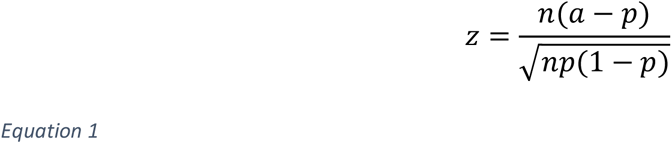

*n* is the number of predictions in the classification problem (i.e., Bernoulli trials), *a* is the fraction of correct predictions, and *p* is the probability of a correct prediction expected by chance, i.e. the reciprocal of the number of classes. Intuitively, this transformation simply expresses the extent to which classification accuracy exceeded chance, scaled by the standard deviation expected by chance. After z-transformation chance level for both classification problems was hence zero.

### Shared response modeling (SRM)

Our hypothesis was that the purely achromatically defined functional retinotopic architecture shared across brains also contained color representations that were equally shared across participants. We therefore used shared response modeling (SRM) (Anderson et al., 2016; Chen et al., 2015; Cohen et al., 2017) to identify the shared functional architecture from responses to the achromatic retinotopic mapping stimulus. For all our SRM analyses we used the implementation that is part of the Brain Imaging Analysis Kit, https://brainiak.org (version 0.7.1). An important advantage of SRM over a related method called hyperalignment (Guntupalli et al., 2016; Haxby et al., 2011) is that it allows for different dimensionalities of individual data matrices (i.e., number of voxels per ROI). This is because SRM estimates linear mappings between ***X****_i_* and ***S*** whereas hyperalignment uses Procrustes transformation to align representational spaces by rotating and scaling vectors between voxel spaces of the same dimensionality. For intelligibility of this manuscript, in the following we briefly describe the mathematical concept of the SRM implementation used.

Let ***X****_i_* be a *v*-by-*d* matrix of fMRI responses (***X**_i_* ∈ ℝ^*v*^ ^×^ ^*d*^) measured for participant *i* with *v* denoting the number of voxels (e.g., within a ROI) and *d* the number of measurements (in our case number of volumes recorded in the retinotopic mapping experiment). In SRM each participant’s response matrix ***X****_i_* is then modeled as a linear transformation of a common response (***S*** ∈ ℝ^*k*^ ^×^ ^*d*^) that is shared by all participants:

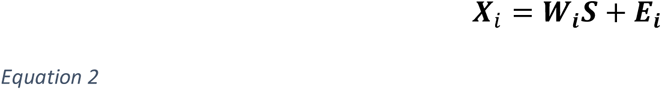

*k* is the number of components in the shared common space. ***W**_i_* ∈ ℝ^*v*^ ^×^ ^*k*^ is the transformation matrix that describes the relationship between the shared responses in the common space ***S*** and each individual’s original response space with matrix ***E**_i_* ∈ ℝ^*v*^ ^×^ ^*d*^ modeling the error between fitted and observed individual data.

SRM estimates the common space matrix ***S*** and individual transformation matrices ***W***_1_, ***W***_2_,…, ***W**_i_*,…, ***W***_*m*_ for each of the *m* participants such that they minimize the Frobenius norm ‖·‖_*F*_ of each error matrix ***E****_i_*summed over all participants:

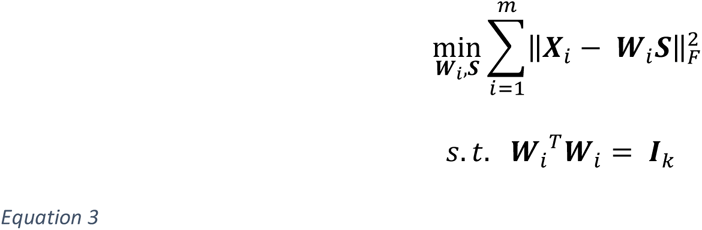

This optimization is performed subject to an orthonormality constraint on each transformation matrix ***W****_i_*. The estimated solutions for ***W****_i_* are therefore similar in interpretation to orthogonality in PCA. Importantly, we can use ***W****_i_* to map patterns of fMRI responses between individual spaces and the common space.

We fit SRMs to data from the retinotopic mapping experiment (Figure 1). This yielded one transformation matrix ***W****_i_* per participant and ROI describing the linear mappings between each individual dataset and the common space of shared responses to purely achromatic, spatially defined stimulation. Let ***D**_i_* ∈ ℝ^*v*^ ^×^ ^216^ be the matrix of *v*-dimensional patterns of responses to the color and luminance stimuli from the main experiment. The transformation matrices ***W**_i_* ∈ ℝ^*v*^ ^×^ ^*k*^ for participant *i* can then be used to map the individual color responses to the *k*-dimensional common space as ***W**_i_*^*T*^***D**_i_*. We repeatedly trained LDA classifiers to distinguish between color (or luminance) categories leaving out one participant’s transformed dataset each time for generalization and then averaging generalization scores across iterations. We obtained confidence intervals for the averaged values from the permuted null distributions (see **Statistical inference** below). The number of common space dimensions *k* for each ROI was chosen by grid-searching a range of 9 candidate values that were evenly spaced on a base-10 logscale from 10^0^ = 1 to 10^2^ = 100 (rounded). Values above 100 dimensions resulted in prohibitively slow model fitting. During cross-validation, each candidate value was applied exclusively to the training set of each iteration and the value resulting in optimal prediction accuracy was used for the final prediction of the withheld test set. The choice of *k* was thus independent from the test set.

**Figure 1.**
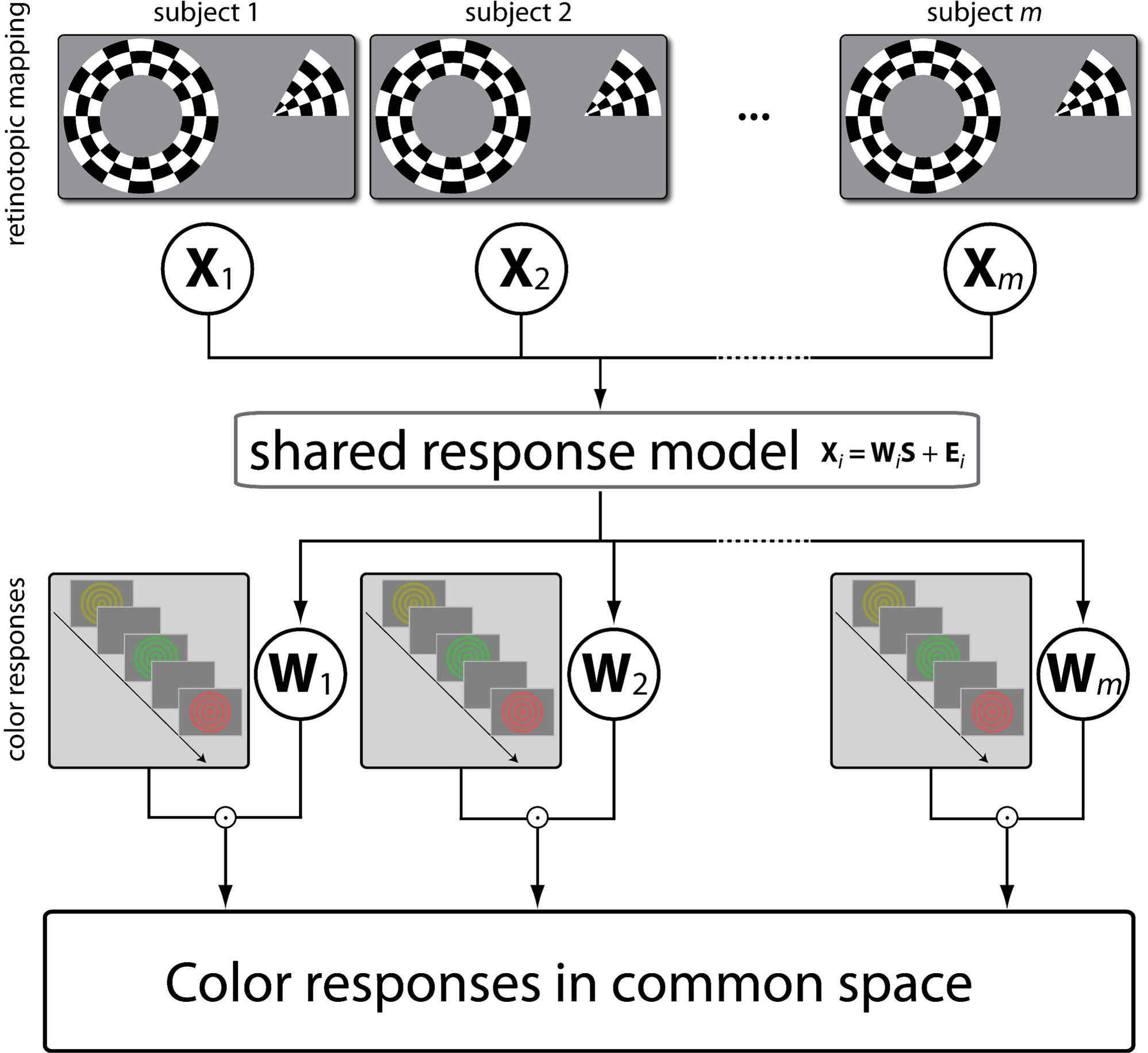
Shared response modeling procedure. Retinotopic mapping data (top) were recorded while participants watched a rotating wedge or an expanding/contracting ring stimulus. Datasets from every participant’s ROI were resampled to the same temporal resolution and entered into a shared response model (Chen2015, Anderson2016). SRM estimates a transformation matrix W_n_ for every participant that maps voxel response spaces from individual ROIs into a 50-dimensional common space (see **Materials and Methods**). Note that only retinotopic mapping data were used to estimate transformation matrices. Color responses (bottom) were measured with fMRI from participants performing a luminance change detection task on red, green, and yellow ring stimuli presented at two luminance levels. Stimuli were shown for 8.5 s with ITI = 1.5 s (see **Materials and Methods**). These fMRI color responses, which were recorded independently from the retinotopic mapping data, were mapped from individual response spaces into the common space using the W_n_ matrices for every ROI and participant.

#### Within-subject preference maps: color & luminance

Since we observed significant classification accuracy of both color and luminance across different brains, we were interested to see whether these representational biases for color and for luminance would be readily visible on the cortical surface of individual brains. This would allow a direct comparison with retinotopic layout of individual visual areas at a single-subject level. If, for instance, there is a preference for a given stimulus class near the visual field meridian, one would expect stronger responses to that class near the visual area boundaries.

We hence estimated the responses to each stimulus class for color and luminance using two separate GLMs in SPM. The time series were modeled in each brain’s individual anatomical space using one predictor for every class and using the same boxcar regressors shifted by 5 s in time (see also **Pattern estimation and preparation**). The resulting maps of stimulus class responses were mapped to the vertices of the individual cortical surface space using FreeSurfer’s estimated registration mapping. For every vertex with a positive (i.e., above-mean) response to any of the stimulus classes, we determined which class was associated with its strongest response. The results were plotted in FreeSurfer’s inflated surface reconstructions with every vertex colored according to the stimulus class with the strongest response.

##### Dependence of between-subject classification on number of shared features

The number of shared features assumed in fitting the SRM determines the dimensionality of common space representations. Note that, in the main analysis (see **Shared response modeling**) the value for this parameter was chosen by grid search using only the training data to learn the optimal value in an unbiased fashion. The BSC analysis was repeated with values for the number of shared features that ranged from 1 to 100 to understand how this affects classification accuracy and how much variance the shared features explain in the data. For each parameter value, we calculated the variance of every voxel time series in the original datasets from the retinotopic mapping session and the main experiment and summed them over voxels and brains. After mapping each participant’s responses from the common space back into original space (using the estimated transformation matrix), the same calculation was conducted on the resulting datasets thereby representing how much variance was explained by the model. Explained variance was quantified as the percentage of this variance relative to the variance computed using the original data. Classification accuracy was calculated as detailed in the paragraph describing the BSC analysis. The 95 % confidence bands are determined as Jeffreys intervals.

#### Dependence of BSC accuracy on number of voxels

Classification accuracies may differ between visual areas because they are different in size and hence contain different numbers of voxels. In order to check to what extent classification accuracies are robust against these differences, we repeated the BSC analysis using the same number of voxels across visual areas. This meant that the number of voxels had to be reduced because the smaller areas impose an upper limit on the maximum number of voxels that can be used for classification. To decide which voxels to select, all voxels were ranked per ROI based on their signal strength in the Fourier analysis of the retinotopic mapping data. The F-value of the test comparing voxel amplitude at the frequency of stimulus presentation to the remaining amplitudes was used as an index for the visual sensitivity of a voxel. All voxels were ranked in a descending order and the top 50, 100, …, 500 voxels (i.e, in steps of 50) were selected. Note that, for any visual area, the smallest ROI across observers limited the largest number of voxels that could be selected from each participant. Thus only areas V1, V2, V3 contained 500 voxels in every participant for example. The number of multiple comparisons that needed to be corrected for accordingly was sometimes lower than in the original BSC procedure. Otherwise it was identical to the main BSC analysis described above.

### SRM searchlight analysis

In addition to ROI analyses, we also applied SRM in a whole-brain analysis to local patterns at every location in the brain normalized to MNI standard space. The purpose of this analysis was to test how specific the shared commonalities were to the exact individual locations of visual areas compared to anatomically aligned patterns of brain responses. Please note that anatomical alignment here refers to the alignment of response patterns which are then subjected to SRM. We do not assume that anatomically aligned voxels already exhibit similar stimulus tunings. Rather, we were interested in determining the difference between modeling shared responses between retinotopically versus anatomically aligned patterns of brain activity.

We used SRM in combination with a searchlight technique that analyzed only local patterns of brain activity within a radius of 3 voxels (“searchlight sphere”) at every location in the whole brain (Kriegeskorte et al., 2006). The number of SRM components *k* was 50 in every searchlight sphere or, if the total number of voxels within the sphere was smaller (e.g. near edges in the brain mask), the number of available voxels. LDA was then used in the same manner as in the BSC ROI analyses to calculate an unbiased estimate of classification performance for every location in the brain, which was then assigned to the center voxel of the sphere. The resulting brain map was then spatially smoothed with a Gaussian kernel of size 6 mm FWHM.

#### Voxel preference prediction across subjects

Here we ask the following question: can we predict color (or luminance) preferences of single voxels of an individual observer based only on the responses to those colors in the remaining brains?

Note that the BSC described above similarly made predictions, but for voxel-response *patterns* – in contrast, here we aim to predict preferences for individual voxels.

This analysis extends previous research showing that the location of category-selective visual regions FFA and PPA could be predicted across brains using a common model space (Haxby et al., 2011) in that it examines the generalizability for more fine-grained visual information than basic level concept distinctions (houses vs. faces).

We pursued this voxel prediction using two approaches:

##### Voxel preference prediction: based on SRM

As in the BSC analysis, SRMs were fit to the retinotopic mapping data of the whole group of brains. The goal is to predict, for each brain’s ROI, the stimulus preferences for all individual voxels in a leave-one-brain-out cross-validation procedure. To this end, the individual mean patterns of responses to every stimulus class from all training brains were mapped into the common space using the individual transformation matrices from the SRM. The transformed patterns were then averaged per stimulus class yielding one group average vector per class. The vector dimensionality depended on the optimal number of shared features as determined by grid-search (see **Shared response modeling**). The group averages were transformed into the individual voxel space of the left-out test brain. The predicted stimulus preference for a voxel was the stimulus class with the largest predicted value, which could then be compared to its true preference based on the main experiment data from that brain, which was used at no point in the prediction pipeline. We refer to this approach as “SRM uni” because it regards preference in terms of which stimulus class causes the maximum univariate response in each voxel. In a variation of this approach referred to as “SRM multi”, we repeated this procedure except that instead of transforming mean vectors of responses to each category, we used the classification weights for each stimulus class estimated using shrinkage LDA. This tested for the possibility that classification-based response patterns may generalize better across brains. Note that also this SRM multi approach predicted preferences for individual voxels.

##### Voxel preference prediction: based on cartesian visual field coordinates

In addition to the above SRM-based approach, we wanted to examine how well a more traditional across-subject voxel-preference prediction approach would fare, and compare this with the above SRM approach. We used phase maps from the polar angle mapping experiment in combination with the phase maps from the eccentricity mapping experiment to obtain the preferred visual field location of every voxel, which we converted to a Cartesian coordinate system with its origin centered on the fovea. A leave-one-brain-out cross-validation procedure was then employed in which all voxels in a given ROI from all but the test brain were assigned a target label based on which stimulus category (e.g., green, red, or yellow) elicited the strongest activation (GLM analysis as described in **Preference maps: color & luminance**). In every cross-validation iteration we fit a *k* nearest-neighbor classifier to the training data, which consisted of as many samples as there were voxels in the ROIs summed over training brains. Each sample consisted of that voxel’s Cartesian x, y visual field coordinates and its target label, which color it preferred. The parameter *k* ranged from 1, 2, …, 9 and was optimized via grid-search using a random split-half cross-validation approach on all training voxels. Nine candidates were considered because there were also nine candidate values for the number of SRM features. The resulting model was cross-validated on the voxels of the left-out test brain by comparing its coordinate-based predicted stimulus preference with its true stimulus preference. Figure 8 shows an illustration of the methods.

Prediction accuracy was quantified using a balanced score metric because the preference for a given color in a ROI’s voxels could not be assumed to be balanced across classes (see `balanced_accuracy_score` in scikit-learn). We tested which of the methods, nearest-neighbor classification without SRM or SRMs using univariate mean responses or classification weights, would do a better job at generalizing across brains (see **Statistical inference** for details).

##### Voxel preference: visualization

Finally, we visualized the dependence of stimulus preference and retinotopic location. The voxel preference predictions from each cross-validation fold were subjected to Gaussian kernel density estimation (KDE, using Scott’s rule to choose bandwidth) to obtain a 2-dimensional density function per stimulus class expressing preference for that class for any visual field location. To obtain a probabilistic interpretation of stimulus preference at any location, we applied a softmax function to the values of each function per stimulus class. This resulted in a probability distribution for every location over the stimulus categories describing how likely it was that a voxel in that location would prefer a given stimulus. We subtracted from each value the probability that a given stimulus class would be preferred under the assumption of a uniform probability distribution, i.e., 1/3 and ½ everywhere for color and luminance, respectively. After converting the probability differences to percentages, a value of x therefore means that the probability of a given class being preferred at this location is increased by x % points. Note that the preference maps are based on predictions that are cross-validated across brains. The prediction of voxel preferences in a given brain therefore requires that any spatial biases in color or luminance processing are reliable across brains used for fitting the model and the brain used for model testing. As an additional check, we performed an odd-even split on the sample of participants and created the same maps based on independent subsets of data to assess split-half reliability. The visualization was performed for the SRM-uni based approach as this obtained the highest scores across ROIs.

### ROI-specificity of SRMs for voxel preference prediction

Since SRMs trained on retinotopic mapping data could successfully be used to predict stimulus preference across different brains, this indicated that they captured to some extent the relationship between retinotopy and color/luminance representation. How specific are these relationships to individual visual areas? Do SRMs trained on different visual areas capture these relationships differently?

To answer these questions, we repeated the voxel preference prediction analysis detailed above using the SRM approach based on the mean patterns of univariate responses to each category. This time, however, we used SRMs that were fit either on data from the same or different ROIs. More precisely, the transformation matrices used to, on the one hand, map individual patterns from individual spaces into the common space, and, on the other hand, to predict stimulus preference in the space of the left-out test brain were based on SRMs fit on data from either the same or different ROIs. All combinations of ROIs were examined for the transformation of the training and test brains and the classification accuracy between congruent (SRMs fit on the same ROI) and incongruent (SRMs fit on different ROIs) were compared. For the incongruent condition, the accuracies were averaged across all combinations where SRMs came from different ROIs. It was tested if the resulting accuracies were significantly larger than chance and if they differed between congruency conditions.

### Statistical inference

#### Within-subject and between-subject classification

All our statistical decisions were based on permutation tests. For WSC we tested the one-tailed null hypothesis that the sample average across all participants’ classification accuracies was equal to or below chance. Since there were three classes in the color classification problem, chance level was *1/3*. Likewise, chance level in the luminance classification was *1/2* as there were two luminance levels.

We created 2000 new label assignments in the following way: For every participant the sequence of training labels was shuffled with the restrictions that labels were only permuted within cross-validation folds, i.e., functional runs, and that permutations were identical across ROIs. Both restrictions were implemented by setting the `groups` and `random_state` arguments in scikit-learn’s `permutation_test_score` accordingly. A group average of classification accuracies was calculated for each ROI and iteration. This way we obtained a null distribution of average classification accuracies expected under the hypothesis that there was no relationship between labels and neural activity patterns.

It was important that permutations were identical across ROIs because they constituted the test family for which we controlled the family wise error (FWE). We formed a new null distribution for all ROIs by taking the maximum of group average classification accuracies across ROIs in every iteration (Nichols & Holmes, 2002). P values were computed as the fraction of permutations that resulted in accuracies that were larger than or equal to the observed classification accuracy. We declared results significant if p was below .05, thereby keeping the type I error probability of falsely rejecting at least one null hypothesis at α = 0.05. For each ROI, the lower and upper limits of the 95 % confidence interval (CI) were calculated parametrically using the standard error of the mean. In the BSC analysis, upper CI limits were obtained by adding the difference between the mean of the (uncorrected) null distribution and its 2.5th percentile to the observed accuracy. Lower CI limits were obtained by subtracting the difference between the 97.5th percentile and the mean from the observed accuracy.

The permutation test for BSC was identical to that for WSC. However, since data from all participants were now combined in the *k*-dimensional functional common space, classifiers were now cross-validated leaving out one participant at a time. Again, labels were permuted only within cross-validation folds, which in this analysis were individual participants.

##### Voxel preference prediction & ROI-specificity analysis

We tested how well different analytical procedures could predict the preference of voxels for different stimulus classes and whether there was a difference in prediction accuracy depending on whether the SRMs used for the training and test brains came from the same or different ROIs (congruent vs. incongruent). In both analyses, it was first tested for each ROI and analysis method whether classification accuracy was significantly larger than chance (1 divided by number of classes) using binomial tests. For pairwise comparisons between analysis methods, we used independent z-tests (Python module statsmodels 0.11.0). The resulting p values for each comparison (against chance and pairwise) were corrected for multiple comparisons across 7 ROIs using Holm-Šidák correction.

#### Retinotopic weight analysis

In this analysis we tested for each ROI whether we could predict from the Cartesian visual field coordinates which color (or luminance level) was preferred by a voxel using a nearest neighbor classifier. Voxel labels were permuted 1000 times, separately within cross-validation folds (participants), and p values were obtained from the null distribution of each ROI. CIs were calculated in the same way as for BSC results. Since there was no correspondence between voxels from different ROIs, we could not control the FWE using the same max statistic approach as in the previous ROI analyses and therefore used Holm-Šidák correction instead to keep α at .05. In order to make sure that the prediction of the highest classification weight was not merely driven by specific pairs of the three colors, we furthermore repeated the same analysis for each pair separately.

#### SRM searchlight analysis

The searchlight analysis yielded a brain map of cross-validated estimates of classification accuracies. In order to test if classification accuracies were significantly larger than what would be expected by chance, we used a one-tailed binomial test instead of a t-test because the classification accuracies from each leave-one-participant-out cross-validation were not independent. The number of correctly classified trials were assumed to come from a sequence of 216 × 15 = 3240 Bernoulli trials with success probability equal to one divided by the number of classes. We applied multiple comparisons correction to the resulting map of p values by keeping the false discovery rate at q = .05 (Benjamini & Hochberg, 1995).

#### Behavioral control analysis

Our classification analysis of neural representations of color assumes that the fMRI data were collected under conditions that differed in color (or luminance) exclusively. In order to rule out the possibility that neural differences were related to differences in task performance between conditions, we therefore conducted a two-way repeated-measures ANOVA of the observers’ reaction times (RT) in the target detection task under the three color and two luminance fixed-effect conditions (3-by-2 design) specifying subject as random factor. A major disadvantage of classical ANOVA is that it only considers empirical evidence *against* the null hypothesis (i.e. indicating that “RTs are different across conditions”) but not against the alternative hypothesis (i.e. indicating that “RTs do not differ across conditions”). Bayesian statistics, however, simultaneously take into account evidence in support of the null *and* alternative hypothesis in a balanced way. Hence we used Bayesian ANOVA (Morey & Rouder, 2018; Rouder et al., 2017) where Bayes factors (BF) quantify the compatibility of the observed data with the alternative hypothesis relative to the null. Specifically, each effect was tested by computing the BF ratio of the globally best fitting model with the best model lacking the effect in question (Rouder et al., 2017). If the globally best model did not include a given effect, the ratio was taken with respect to the best model that did include the effect in question (resulting in BFs favoring the null). As an example, a BF of 3 means that the data are three times more likely to have occurred under the alternative hypothesis than the null. Fixed-effect priors were specified for “medium” sizes by applying the default *r* scale of .5. Normality assumptions were checked with Shapiro-Wilk’s tests and by visual inspection of Q-Q plots of the residuals (not shown). Sphericity was tested using Mauchly’s test.

#### Retinotopic binning analysis

The present paper shows results from a novel analysis testing for a dependency between retinotopic properties of fMRI voxels and their selectivity for different colors. By identifying the shared responses to visual stimulation by both color and purely retinotopically defined input, it allowed us to visualize this dependency by mapping the classification weights of the shared responses to visual field maps.

Given its novelty, we assured ourselves of the validity of our conclusions from our analysis by complementing it with a more conventional procedure that had been applied previously to test for retinotopic biases (Beckett et al., 2012; Freeman et al., 2011; Larsson et al., 2017) and for which our conclusions – if true – make testable predictions as well. In this procedure WSC of color is conducted as follows: all voxels in a given ROI are binned into a certain number of bins and the voxel responses to the colors are averaged per bin. The dimensionality of each response pattern in a ROI containing *v* voxels binned into *b* bins is thus reduced from *v* to *b*. Now, to test for a retinotopic bias in this scenario, binning is carried out in two different ways. In one analysis, the phase in which voxels respond to the retinotopic mapping stimulus (polar angle or eccentricity) and which ranges from 0 to 2π is used to assign voxels into bins consisting only of voxels with similar phases. This preserves some aspect of the retinotopic structure in the data. In the control analysis, voxels are binned randomly. If classification is mediated at least in part by retinotopic bias, one would expect that retinotopy preserving binning yields higher classification accuracy than random binning.

The two analyses are conducted for several values of *b* chosen from a base-10 logarithmic scale ranging from 0 to 3 in steps of .2: *b* ∈ [10^0^, 10^.2^, …, 10^*b*^, …, 10^3^]. The analysis was carried out separately for every participant and ROI. If *b* exceeded the number of voxels in any of the participants in a given ROI, this binning step was skipped. We statistically compared retinotopic and random binning with paired t-tests corrected per ROI for multiple comparisons across 15 binning steps (or fewer, see above) using the Holm-Šidák procedure.

## Results

The goals of the experiment were to examine whether representations of chromatic signals were shared with a spatial neural code, whether spatial color specificity was shared across human brains, and whether this was area specific. For this reason, we used brain responses to achromatic, spatial stimuli to construct a neural space that was shared between observers (Figure 1). We then tested whether this neural space generalized to decode color responses in specific, individual brains from color responses in other observers’ brains -- in other words -- to predict color across brains. We then tested whether an area-agnostic common space across participants allows predicting colors only within a given area or across areas.

### Within-subject classification of color and luminance (WSC)

Decoding color across different brains requires that color can be decoded within one and the same individual brain. For this reason, we first performed pattern classification analyses on fMRI activity from ROIs at the level of individual subjects. The same logic applied to luminance decoding. Figure 2 shows classification accuracies z-transformed (n = 216) individually and then averaged across participants (n = 15). Color as well as luminance could be decoded significantly better than chance in all ROIs under investigation (permutation tests, each p < .01, FWE-corrected for seven ROIs). For color (chance level: 33 %), the following classification accuracies applied for each ROI, averaged across participants. V1: 57 % (z = 7.38, 95 % CI [6.51, 8.25]), V2: 55.4 % (z = 6.89, 95 % CI [5.94, 7.84]), V3: 52.84 % (z = 6.08, 95 % CI [5.13, 7.03]), hV4: 51.2 % (z = 5.56, 95 % CI [4.21, 6.91]), VO1: 46.3 % (z = 4.03, 95 % CI [3.01, 5.05]), LO1: 44.8 % (z = 3.56, 95 % CI [2.66, 4.46]), LO2: 42.5 % (z = 2.86, 95 % CI [1.72, 4]). For luminance (chance level: 50 %), classification accuracies were as follows. V1: 61.9 % (z = 3.51, 95 % CI [2.32, 4.7]), V2: 65 % (z = 4.41, 95 % CI [2.81, 6.01]), V3: 66.7 % (z = 4.92, 95 % CI [3.45, 6.38]), hV4: 59.1 % (z = 2.67, 95 % CI [1.84, 3.5]), VO1: 56.7 % (z = 1.98, 95 % CI [1.16, 2.8]), LO1: 63.5 % (z = 3.96, 95 % CI [2.71, 5.22]), LO2: 59.9 % (z = 2.9, 95 % CI [1.69, 4.11]). Individual response patterns in every ROI thus were reliable enough across fMRI runs to linearly predict color and luminance condition.

**Figure 2.**
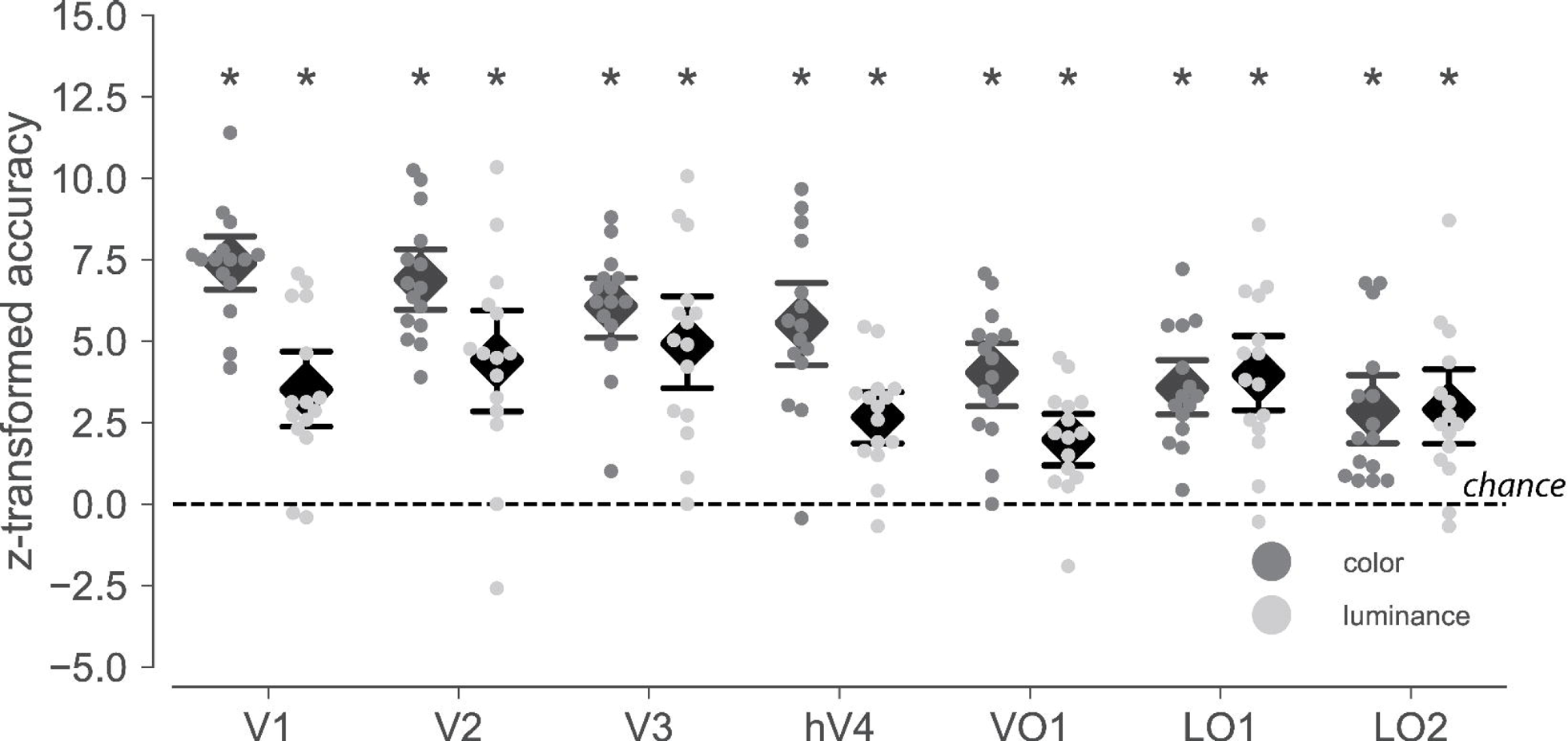
Multivoxel pattern classification results for within-subject classification. For every ROI and participant, cross-validated (leave-one-run-out) classification accuracies were obtained for the prediction of color (dark dots) and luminance (light dots). Accuracies are transformed to z-values using the normal approximation to the binomial distribution (n = 216) so that both classifications share the same y-axis and accuracies expected by chance is 0 in both cases. Black diamonds indicate mean accuracy averaged across participants and error bars represent the parametric 95 % CI. Asterisks mark accuracies with p values below .01 (permutation tests, 2000 iterations, FWE corrected across 7 ROIs). As shown, classification accuracies were significantly larger than chance for both color and luminance classification.

### Between-subject classification of color and luminance (BSC)

Next, we quantified how consistent chromatic neural processing was across individual brains within a neural space that was shared across subjects and defined by responses elicited by a solely achromatic and spatially defined retinotopic mapping stimulus. Individual color responses were mapped from independent measurements to that shared response space, and the classifiers were trained to predict the color (or luminance) of the stimulus a person was seeing. In contrast to WSC, fitted classification models were cross-validated across participants instead of runs. As can be seen in Figure 3, accuracies for luminance and color classification across different brains significantly exceeded chance in areas V1-V3. Additionally, color could be decoded across participants significantly better than chance in areas hV4 and LO1 (permutation tests, p < .01, FWE-corrected for seven ROIs). For each ROI, significant BSC accuracies for color were as follows. V1: 44.7 % (z = 13.7, 95 % CI [8.23, 19.0]), V2: 39.8 % (z = 7.75, 95 % CI [4.23, 11.09]), V3: 39.6 % (z = 7.57, 95 % CI [4.11, 10.75]), hV4: 39.5 % (z = 7.42, 95 % CI [3.27, 11.47]), LO1: 38.8 % (z = 6.6, 95 % CI [2.1, 10.9]). Accuracy in VO1 was 34.0 % (z = 0.78, 95 % CI [−2.11, 3.6], p = .873, FWE corrected) and thus not significant. In LO2 accuracy was 36.0 % but did not survive FWE correction (p = .021, uncorrected): z = 3.17, 95 % CI [.14, 6.14], p = .285, FWE corrected. For luminance, significant classification accuracies were as follows. V1: 56.2 % (z = 7.03, 95 % CI [2.9, 11.36]), V2: 56.4 % (z = 7.24, 95 % CI [4.23, 11.09]), V3: 55.2 % (z = 5.94, 95 % CI [22.22, 9.46]). This suggests that measurements of how purely spatially defined stimulation activates the brains of different individuals is sufficient to predict colors and luminance from brain activity in any one of them using a classifier trained on data from the remaining participants.

**Figure 3.**
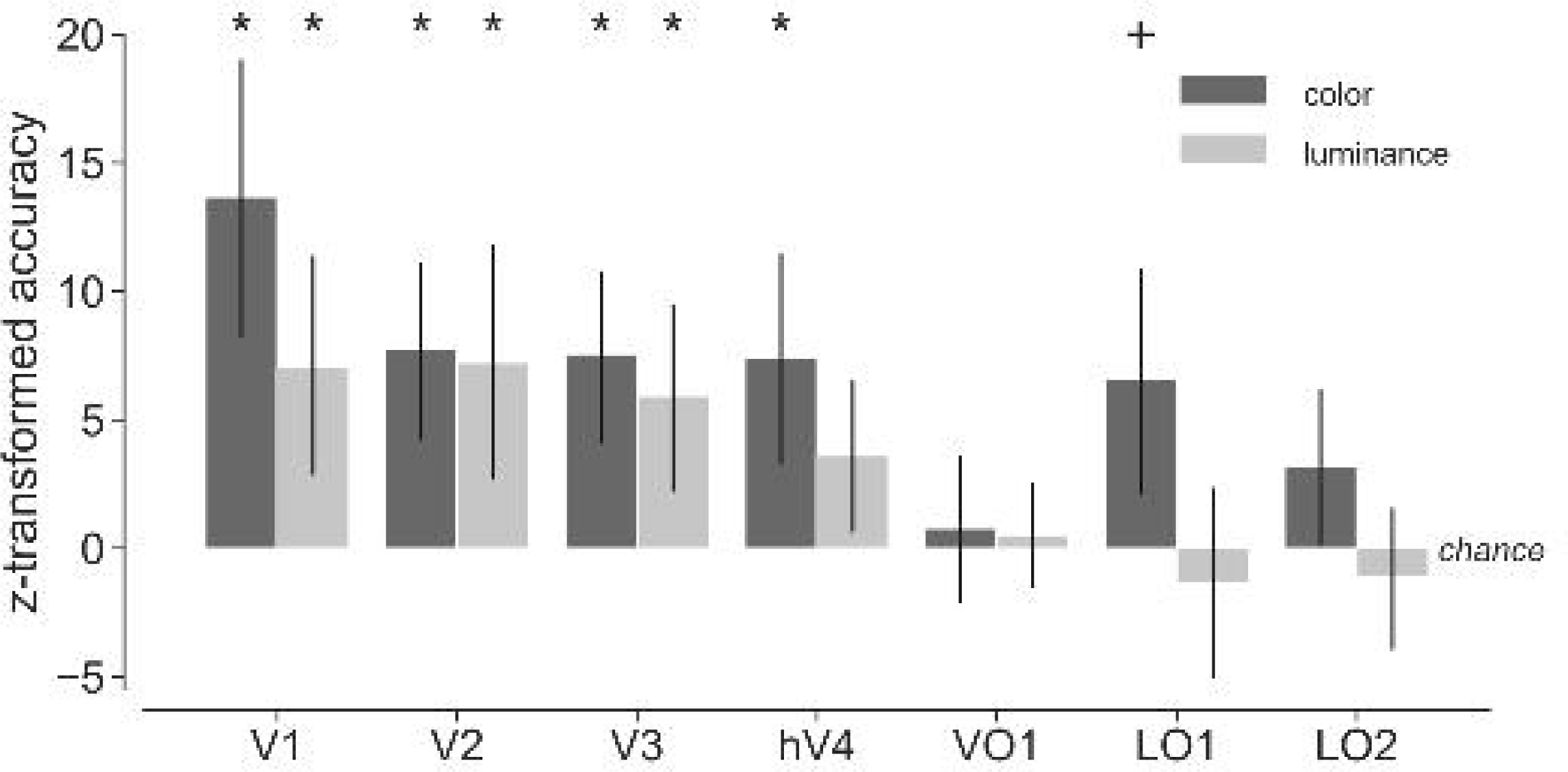
Multivoxel pattern classification results for between-subject classification. Individual responses were transformed to the common space. Transformation matrices were estimated using the shared response model fit to the independent set of retinotopic mapping data. Classification accuracies were cross-validated by leaving out the participant during classifier training whose data were used for testing (leave-one-subject-out). The normal approximation to the binomial distribution was used to convert accuracies to z-values (n = 3240). Note that prediction accuracies are represented as bars rather than individual dots (see Figure 2) because the prediction accuracies for individual participants are no longer independent. Error bars denote the 2.5th and the 97.5th percentile of the permuted null distributions. Asterisks and plus sign denote accuracies significantly exceeding chance at p < .01 and p < .05, respectively (permutation tests, 2000 iterations, FWE corrected across 7 ROIs). Both color and luminance could be predicted across subjects from areas V1-V3. Additionally, color could be predicted from hV4 and LO1.

#### Individual preference maps: color and luminance

To check for the possibility that any retinotopic biases might be readily visible from the inspection of the response maps on the cortical surface reconstructions, we transformed the mean responses to every stimulus class to surface space and determined which of them caused the strongest response at each cortical location. Figure 4 shows the results of this analysis for the classification of color. At the individual level, no systematic relationship between cortical location and color that is consistent across brains is apparent from the maps. Figure 5 shows the same maps for the classification of luminance. Preference for high intensity luminance is shown in yellow, preference for low intensity luminance is shown in blue. Likewise, no consistent relationship between preference and location can be observed. The dependence of color representation upon cortical location thus appears to be too subtle to be apparent by visual inspection from individual brains.

**Figure 4.**
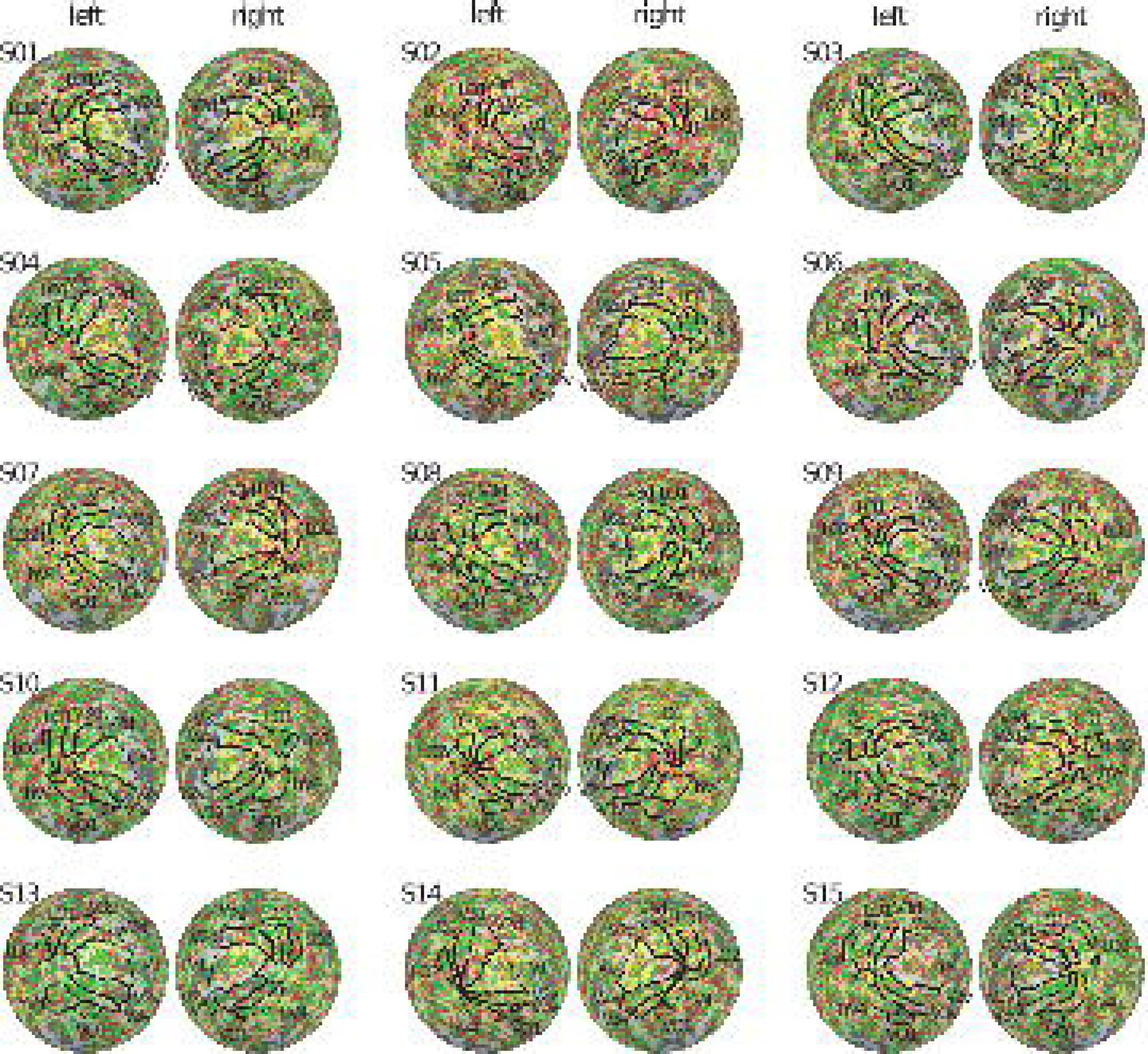
Color preference maps. To check if any retinotopic response biases were already evident from the pattern of univariate responses, the univariate voxel GLM coefficients for each color category were projected onto each participant’s inflated cortical surfaces of the left and right hemispheres. Each vertex was painted in the color of a given stimulus if its GLM coefficient was positive and larger than the coefficients for all remaining colors. There is no obvious correspondence between preference maps across participants.

**Figure 5.**
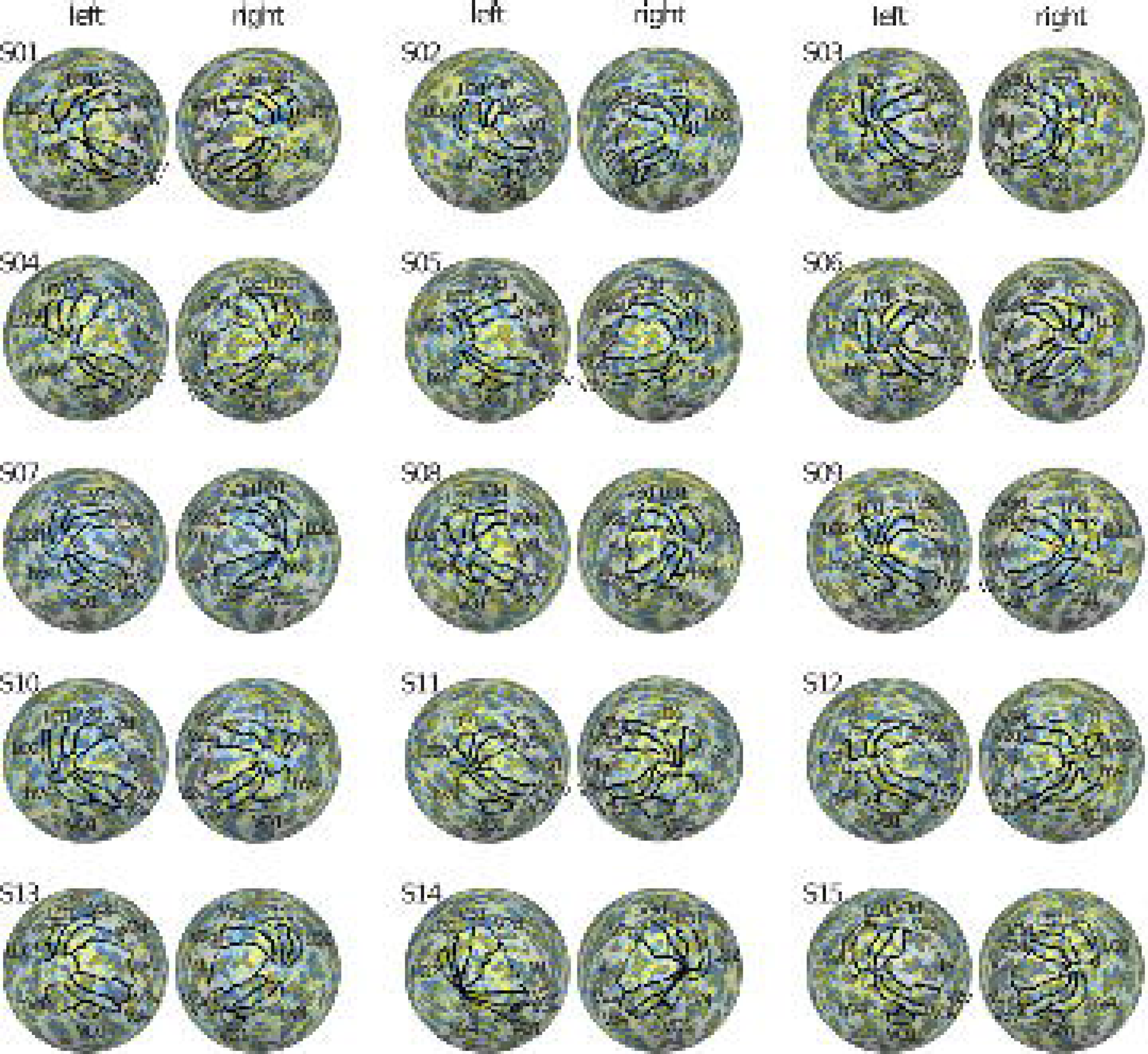
Luminance preference maps. Same as Figure 4 except that surface maps here show vertex preference for high luminance in yellow and for low luminance in blue. Again, there is no obvious correspondence between preference maps across participants.

##### Dependence of between-subject classification on number of shared features

Shared response modeling requires a parameter that controls the dimensionality of the shared common space. To understand how much variance needs to be explained by the model to enable above-chance BSC accuracy, we repeated the analysis using different numbers of shared components between 1 and 100. Figure 6 shows results for the classification of color (top panel) and luminance (bottom panel). The blue curves show how much variance is explained by the SRM components as a function of number of features, for retinotopic mapping data (solid lines) and for data of the main experiment (dashed lines). The former are generally higher than the latter because the model features are determined based on the retinotopic mapping data. More components naturally explain more variance in the training set. This also applies to the data from the main experiment, which were not used to fit the components. Classification accuracies (orange lines) increase with the number of shared features but plateau or decrease slightly from typically between 10 to 20 features onwards. The peak of classification (orange) hence occurs approximately where 15 to 30 % variance are explained in the training data (blue curve). This is also where a weakly pronounced “elbow” in the blue curves is visible. The results indicate that using more features does not necessarily lead to better classification accuracy and that accuracy peaks are typically found with low numbers of features that explain between 15 and 30 % of the variance in the training set. The dependence of classification accuracy on the number of features emphasizes the need for an unbiased selection procedure of this parameter like the one we used in the main analysis.

**Figure 6.**
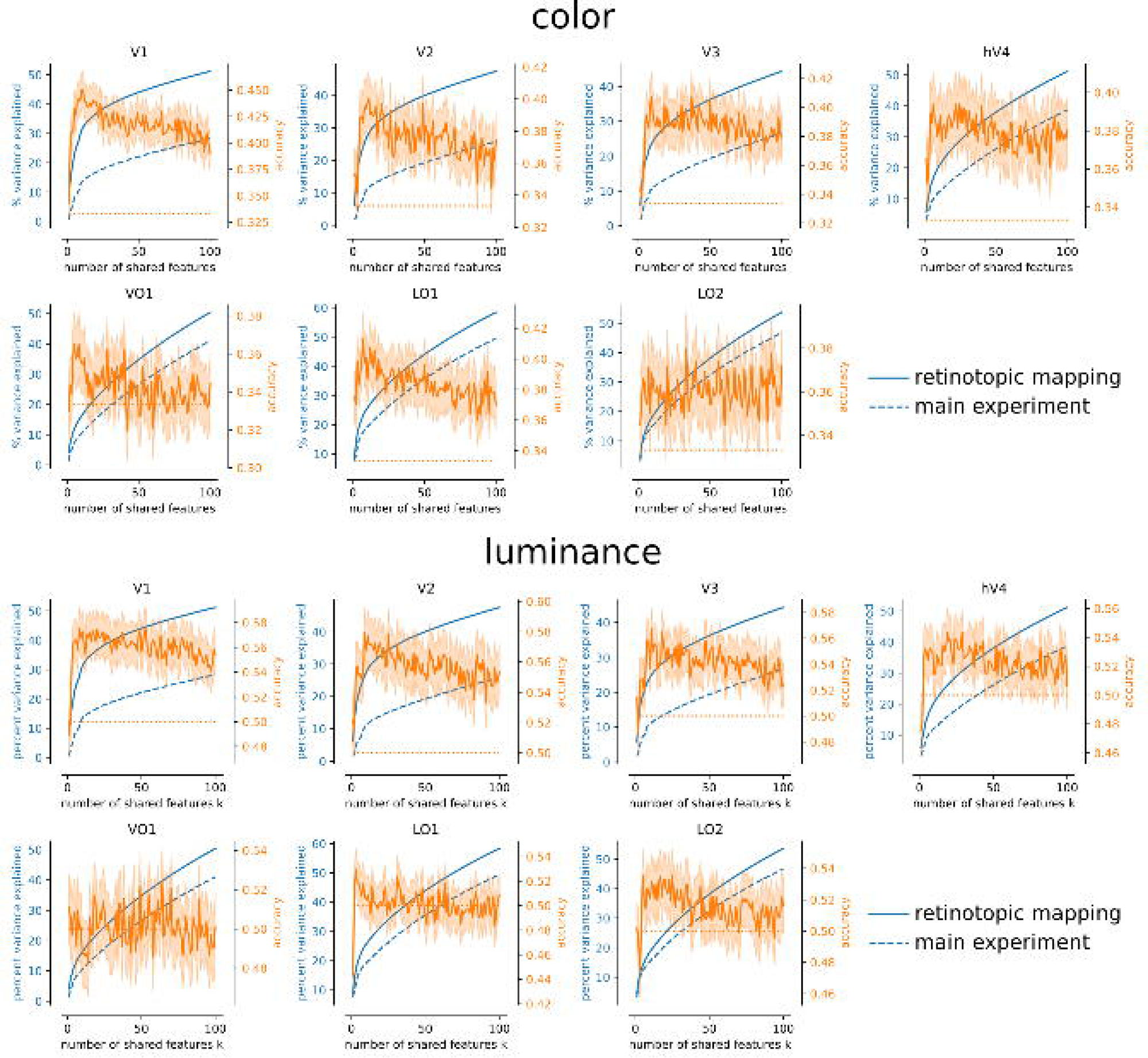
Dependence of BSC accuracy on number of shared features. **Top:** explained variance and decoding accuracy for the color data. **Bottom:** same for luminance data. The x-axis denotes the number features used for the SRMs’ fit to the retinotopic mapping data. The left y-axis (blue) denotes the percent variance explained by the features of the fitted models for the retinotopic mapping dataset (solid blue lines) and the color (luminance) responses from the main experiment (dashed blue lines). The right y-axis (orange) denotes the accuracy of the between-subject classification of color (luminance) and its 95 % CI (Jeffreys interval) (solid orange, see also Figure 3). Chance level is indicated by the dotted orange line. A higher number of shared features resulted in more explained variance for the retinotopic dataset. This applies to the main experiment as well although it was not used for model fitting. Classification accuracy only increased with more shared features for low values but shows a downward trend beyond approximately 10 to 20 features. This turning point coincides with a weakly pronounced “elbow” in the explained variance curves. Few shared features are thus sufficient to capture the information shared across observers. Note that the number of shared features in the main analysis (Figure 3) was selected by grid-search on the training set.

##### Dependence of BSC accuracy on number of voxels

We examined to which extent the results from the BSC analysis were robust against changes in the number of voxels used for classification. That way the number of voxels could be held equal across ROIs, which otherwise typically differ due to differences in size between visual areas. Figure 7 shows the results for BSC of color (top) and luminance (bottom). Descriptively, classification accuracies tend to be lower compared to the results from the original BSC analysis (Figure 3), as can be expected given the lower number of voxels. Classification accuracy in V1 is fairly robust in that it is significantly larger than chance across all numbers of voxels tested. In the majority of selection steps classification accuracy was likewise significantly larger than chance in hV4 (4 out of 7) and LO1 (6 out of 7). Classification accuracy was more dependent on the number of voxels in V2 and V3. It exceeded chance level at a more lenient uncorrected significance threshold (V2: 6 out of 10; and V3: 5 out of 10). Accuracy was not above chance at any selection step in VO1 and in LO2 only at one step (uncorrected). BSC accuracy was thus largely robust across selection levels in V1, hV4, and LO1. In V2 and V3 above-chance classification was also possible with fewer voxels but it was more contingent on the number of voxels used for the analysis.

**Figure 7.**
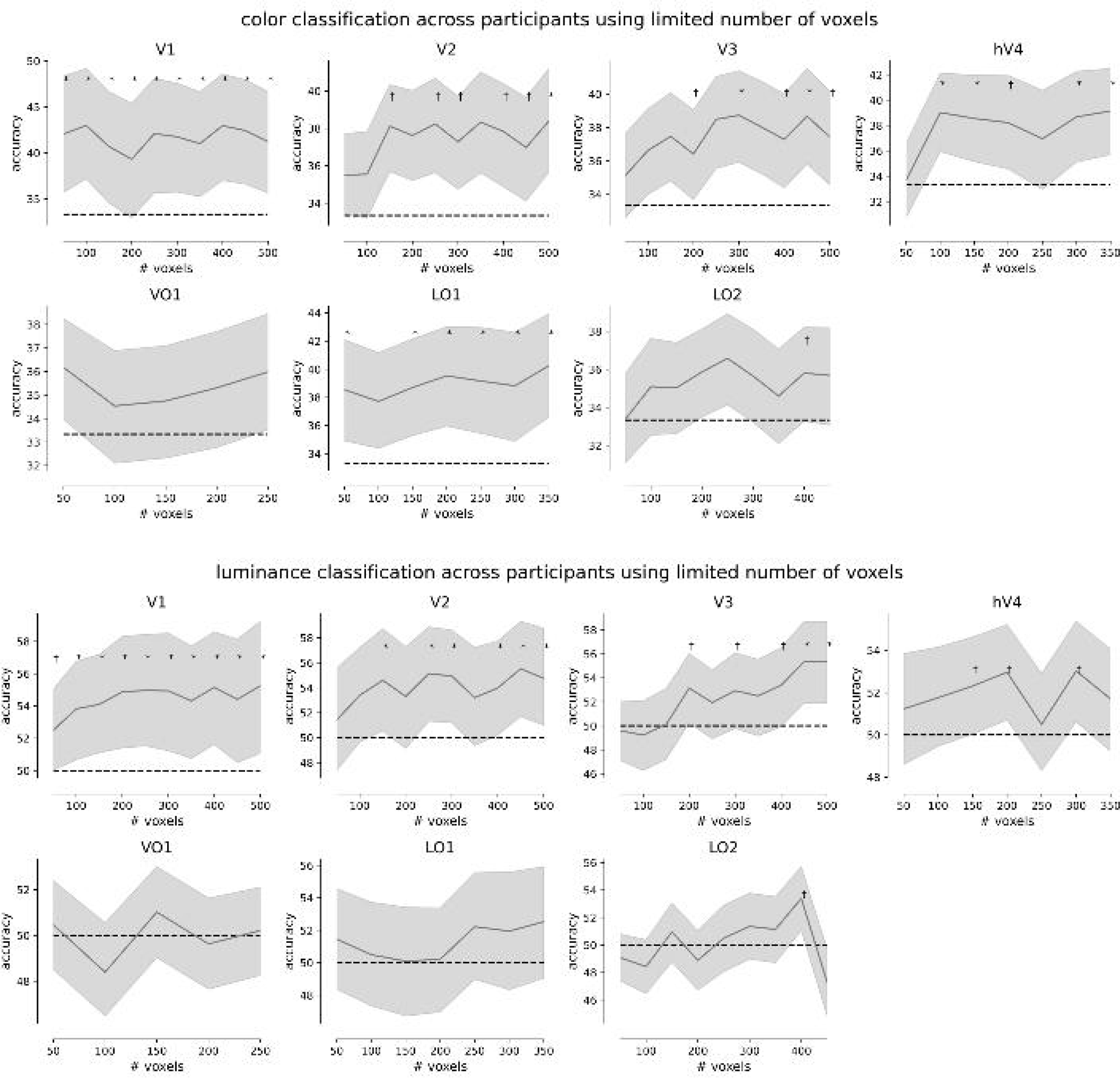
Dependence of BSC accuracy on number of voxels. Since ROIs differ in terms of the number of voxels they contain, an analysis was conducted with different numbers of voxels. Voxels within ROIs were ranked according to signal quality (F value from the Fourier-based analysis of the retinotopic mapping data) and only the best few voxels were included in the analysis. Curves represent classification accuracies as the mean percent correct classifications across the group and confidence bands are the 95 % CI. Asterisks mark accuracies that are significantly larger than chance (p < .05, Holm-Šidák corrected for the number of ROIs, which could be smaller than seven because some ROIs did not contain enough voxels in all tested subjects) and daggers denote p < .05 (uncorrected).

As for luminance, classification accuracy in V1 similarly exceeds chance significantly at 9 out of 10 selection steps. In V2 it is significantly above chance in 6 out of 10. V3 showed less robustness with accuracy surpassing a more lenient uncorrected significance threshold at 5 out of 10 steps. In hV4 significant classification was observed at 3 out of 7 and in LO2 at 1 out of 9 voxel selection steps (uncorrected). Decoding thus was robust across selection steps in V1 and V2, whereas V3 decoding required larger numbers of voxels.

### Voxel preference prediction: large-scale retinotopic color biases

Since our BSC results were based on shared responses to retinotopic mapping experiments, we were interested in a closer examination of any large-scale retinotopic biases mediating across-subject color decoding. We therefore tested how accurately we could predict to which color (or to which luminance level) each voxel in every ROI of a given test brain would respond most strongly, i.e., which it would prefer. Importantly, this prediction would be based only on the retinotopic mapping data and on color/luminance data from brains that were independent from the test brain. We compared different methods, which are illustrated in Figure 8 A and B. The first relied on SRMs, the second on traditional cartesian retinotopic projections combined with kNN classification.

**Figure 8.**
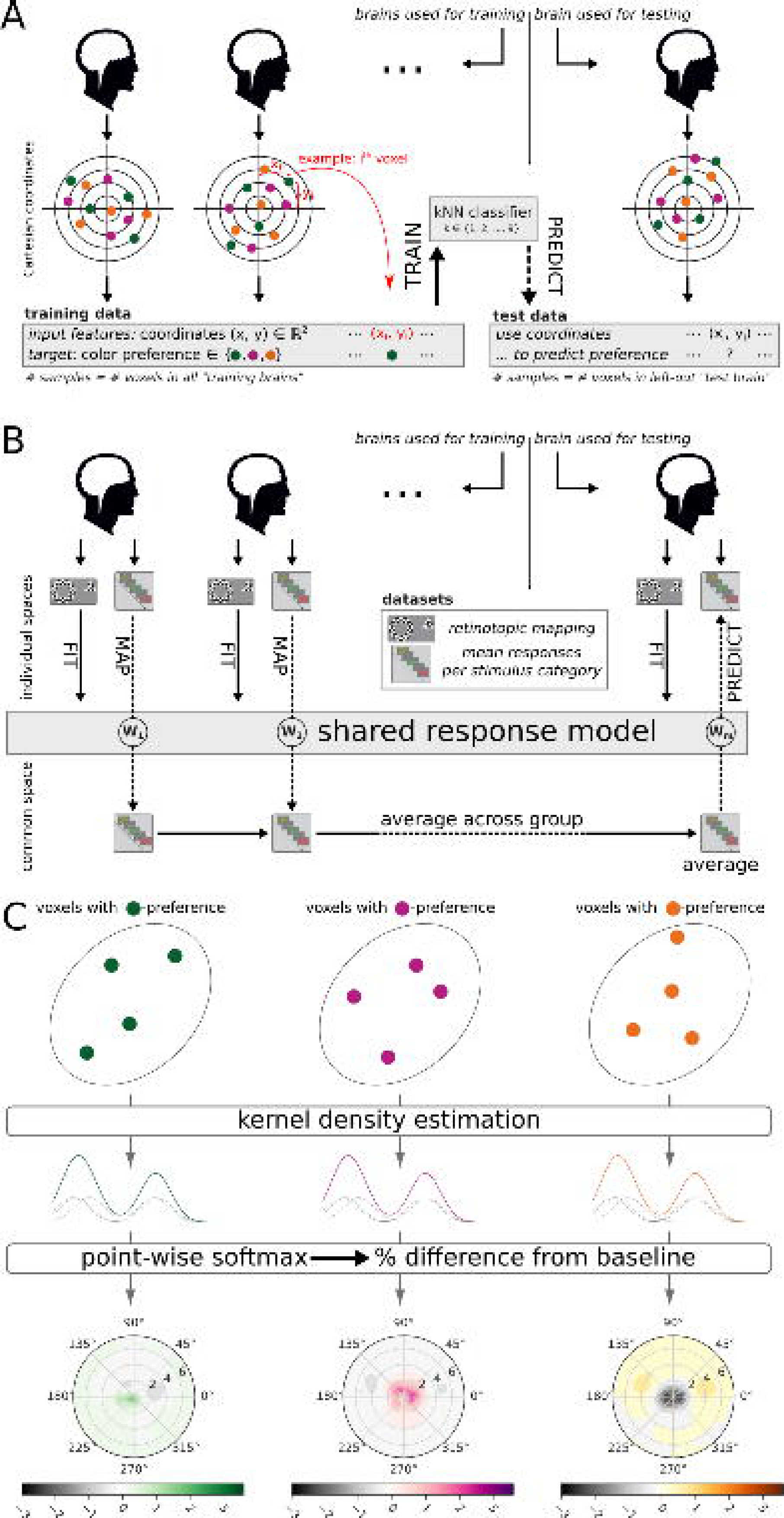
Preference prediction analysis. Illustration of two methods to predict stimulus preferences (color or luminance) of individual voxels within a ROI in an independent, withheld brain based on activity observed in the remaining brains. **(A)** Prediction based on traditional cartesian retinotopic voxel locations coupled to kNN response preference classification. All voxels from a given ROI were extracted from all but one observer’s brains (leave-one-subject-out cross-validation). Each voxel was labeled according to which of the stimulus category (e.g., green, red, or yellow) elicited its strongest response, i.e., its target variable. A nearest-neighbor classifier was trained to predict the target from the Cartesian retinotopic coordinates of that voxel. The classifier was then cross-validated on the left-out brain where the predicted voxel preferences based on the retinotopic coordinates were compared to the true voxel preferences. **(B)** Prediction based on SRM (common space). SRM provides direct mappings between individual and common space representations of brain activity. The retinotopic mapping data were used to obtain a shared response model for the whole group of participants. The **W** matrices obtained from the SRM were used to map every vector of voxel responses for each stimulus category from all individual ROI voxel spaces to the common space. The vectors in common space were averaged across participants for each category and the result was mapped into the individual ROI voxel space of the left-out brain. The preference prediction for each voxel in the test brain was determined as the stimulus category with the strongest predicted response. In a second version of this model, instead of the mean univariate responses to each stimulus category, the multivariate classification weights for each stimulus category were used for prediction. **(C)** Visualization of predictions. To visualize the stimulus preference across retinotopic space, we estimated the preference for each visual field location from the predicted voxel preferences using kernel density estimation (KDE). This yielded one 2D density function per stimulus category. At each visual field location, a softmax function was applied to the different function values from each category to obtain a probabilistic interpretation in percent of stimulus preference at that location. Finally, the baseline percentage (100 % / number of categories) was subtracted from each value to facilitate the distinction between positive and negative deviations from uniformity.

As Figure 9 (left) shows, **color preference** could be predicted significantly better than chance using several of the approaches tested. Importantly this means that individual areas do indeed have large scale, retinotopically mapped color biases, and that these can robustly be revealed using distinct approaches. In more detail, we found that the number of ROIs where accuracy was significantly above chance was largest for the SRM-uni approach (5 out of 7) compared to “SRM multi” (2 out of 7) and the nearest neighbor approach (3 out of 7). The five ROIs significant in SRM-uni were the same as those where significant classification of color across brains (BSC) was observed (all p < .01): V1 (38.4 %, 95 % CI [37.8 %, 39 %]), V2 (36 %, 95 % CI [35.4 %, 36.6 %]), V3 (36.9 %, 95 % CI [36.2 %, 37.5 %]), hV4 (35.3 %, 95 % CI [34.4 %, 36.2 %]), and LO1 (37.7 %, 95 % CI [36.7 %, 38.7 %]). There was no ROI where the “SRM uni” accuracy was significantly surpassed by either SRM-multi or the nearest neighbor approach. In contrast, “SRM uni” accuracy was significantly larger than the two other methods in LO1 (nearest neighbor: p < .05, SRM-multi: p < .01), V3 (both p < .01). In V1 it exceeded that of SRM-multi (nearest neighbor: p = .2392, SRM-multi: p < .01) and in V2 that of the nearest neighbor approach (nearest neighbor: p < .01, SRM-multi: p = .249). In hV4 SRM-uni was the only method to show accuracy that was significantly above chance (p < .01, nearest neighbor: p = .3383, SRM-multi: p = .2407). In sum, while color preference could be predicted using different methods, the SRM approach based on mass univariate patterns of color responses yielded the highest generalization accuracy across brains.

**Figure 9.**
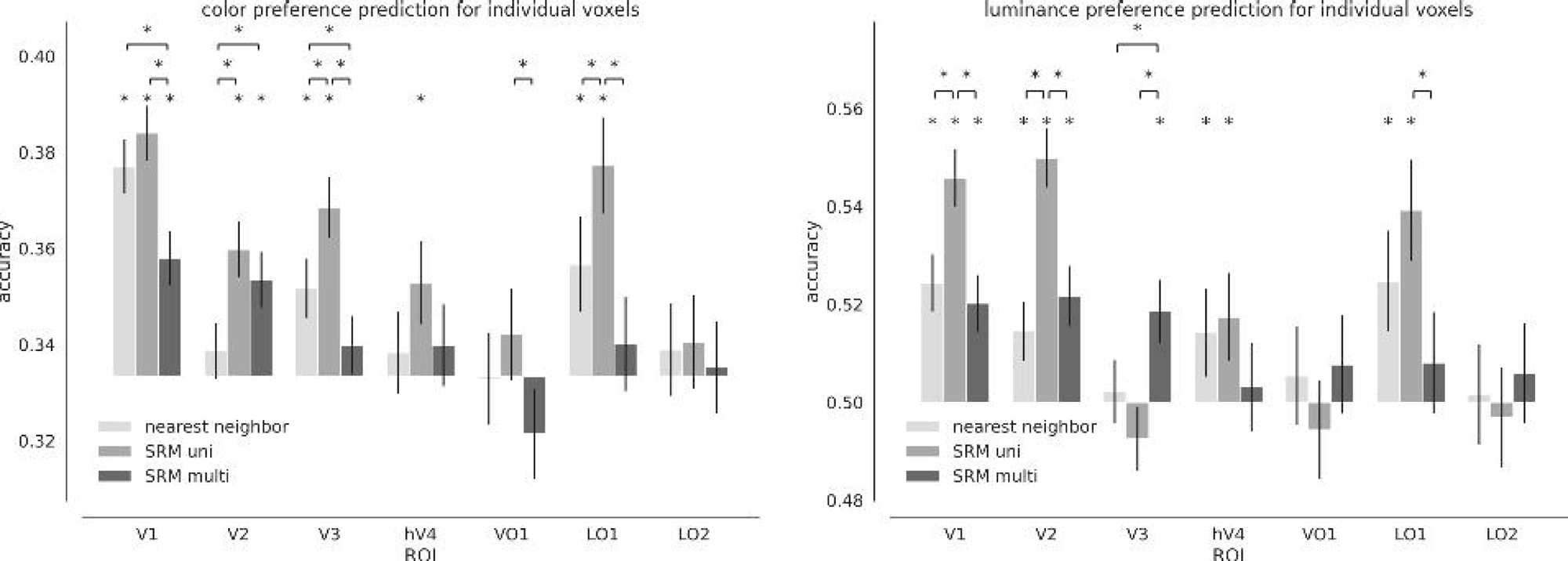
Preference prediction results. Comparison of the two procedures with and one without SRM to predict voxel preferences across different brains for color (left) and luminance (right). Accuracy is the fraction of correctly classified voxels across all brains and error bars represent 95 % CIs. Binomial tests were used to test if preference prediction accuracy was significantly larger than chance. Pair-wise comparisons were made using independent z-tests. Asterisks denote significance at p < .05, Holm-Šidák corrected for seven ROIs. While different methods could be used to predict stimulus preference with above-chance accuracy, the SRM approach with univariate measures of voxel preference frequently outperformed its alternatives.

As for **luminance classification** (Figure 9, right), both SRM-uni and nearest neighbor approaches yielded accuracies that were significantly above chance in the same four ROIs, V1, V2, hV4, LO1 (all p < .01). SRM-multi accuracies were significantly better than chance only in V1, V2, and V3 (all p < .01). However, SRM-uni outperformed both other approaches in V1 and V2 and SRM-multi in LO1 but neither of them in hV4. Only in V3 SRM-multi was the only method with above-chance classification accuracy, which exceeded those of the remaining approaches. Although there was good agreement between all methods in V1 and V2, only SRM-multi showed preference prediction in V3 whereas the other two methods instead showed significant accuracies in hV4 and LO1. With the exception of V3, SRM-uni, however, showed the highest accuracies at generalizing luminance preference across brains.

Since across-subject commonalities in retinotopic voxel preferences were detected most robustly with SRM-uni, we proceeded with this method to map stimulus preference across the visual field.

#### Stimulus preference across the visual field

Given the above statistical evidence for large scale, retinotopically mapped color biases in individual visual areas, we aimed to visualize them as illustrated in figure 8C. We visualized the predictions that the SRMs made for individual voxels at different locations across the visual field by applying KDE on the cross-validated predictions made for each brain (see Methods: Voxel preference prediction across subjects) and converting them to probabilities using a softmax operation.

Color preference maps can be seen in Figure 10. The maps in the large polar plots are based on the whole sample of brains. Note that these maps are based on models that were fit to and evaluated on independent datasets and that successful prediction of voxel preferences presupposes that regional biases are reliable across datasets. The small plots serve as an additional assessment of reliability. They are based only on data from the two halves of the sample after an odd-even split and therefore visualize split-half reliability. Descriptively, the results from the whole sample showed a systematic relationship between visual field location and color preference. In V3 for instance voxels representing parafoveal locations showed a preference for yellow. At intermediate eccentricities along the horizontal meridian the analysis revealed a preference for green. More peripherally, our analysis showed a preference for red in the upper and lower visual fields. Visual inspection of the pairs of maps resulting from the odd-even split show good agreement in the ROIs where above-chance BSC was observed (Figure 3), indicating sufficient reliability.

**Figure 10.**
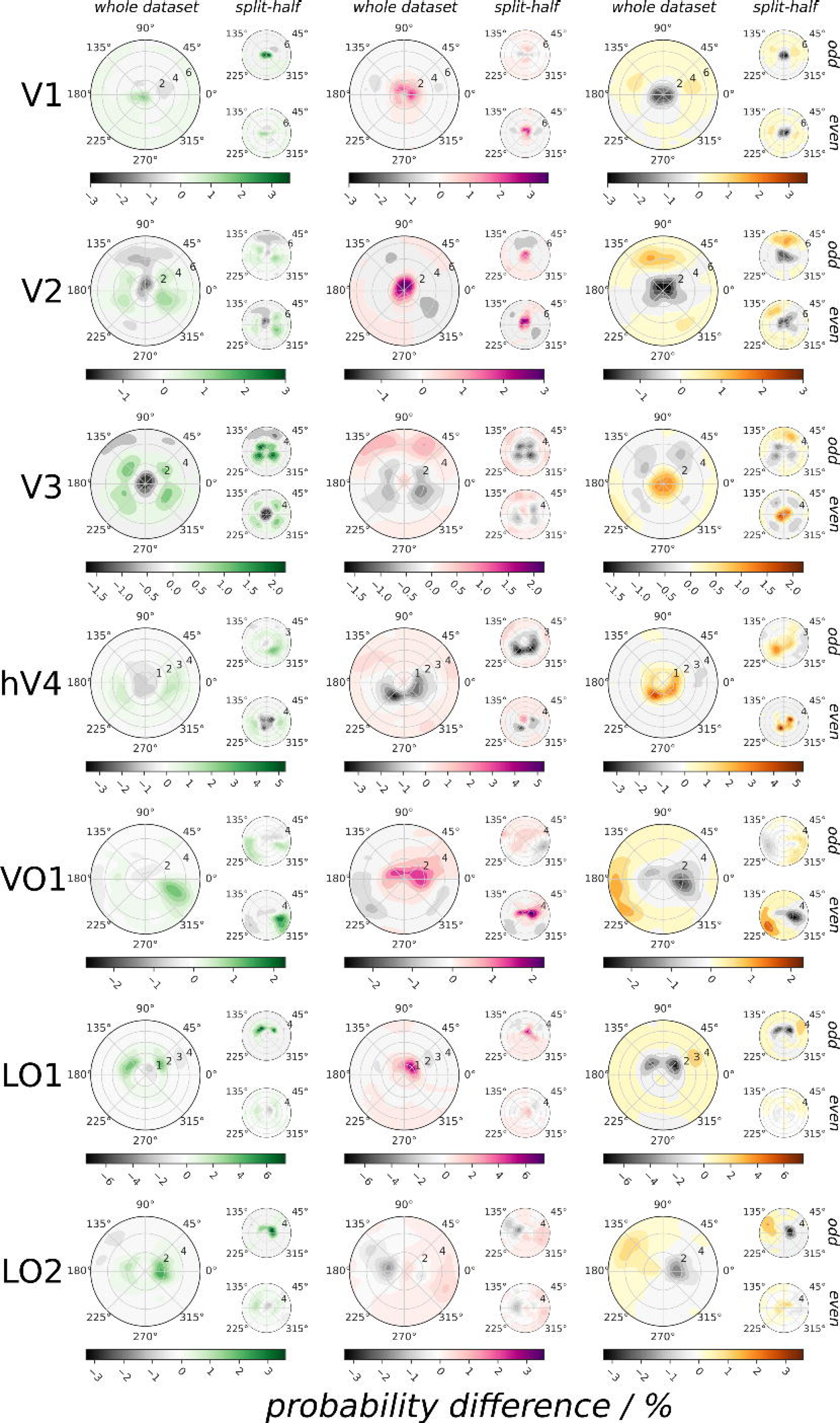
Color preference across the visual field. Color preference was visualized as explained in Figure 8 **(C)**. Each of the three columns indicate the probability of a voxel to prefer the green, red, yellow stimulus for every visual field location. SRM-uni based maps are shown as it achieved highest scores (see Figure 9). The plotted values are the difference between this probability and the “baseline probability”, i.e., the probability that would be expected if it were distributed uniformly across the three stimulus categories, i.e., 33.3 %. Positive (colored) values thus represent regions where the preference probability for a given color is larger than baseline, negative (gray) values represent regions where it is lower. Large plots show results for analysis using all participants’ datasets, small plots show results using only subsets of the data based on an odd-even split. Visual areas show characteristic dependencies between retinotopic location and color preference: V1 for instance exhibits a preference for green or red near the fovea and less for yellow while the opposite is true at higher eccentricities.

Importantly, the maps differed between ROIs. For example, whereas in V3 the yellow stimulus was preferred at parafoveal locations, this was not the case in V1, where preference for yellow was more pronounced at more peripheral locations. These results speak against inheritance of large-scale color-biases along the processing hierarchy and for area-specific properties, which will be examined in more detail in the following section.

Figure 11 shows the same results for luminance. Preference maps again differed across brain regions: While in V1 and V2 the preference for stimulation at high luminance decreased with increasing eccentricity for instance, the preference for high luminance stimulation in V3 was more pronounced in the lower visual field than in the upper visual field.

**Figure 11.**
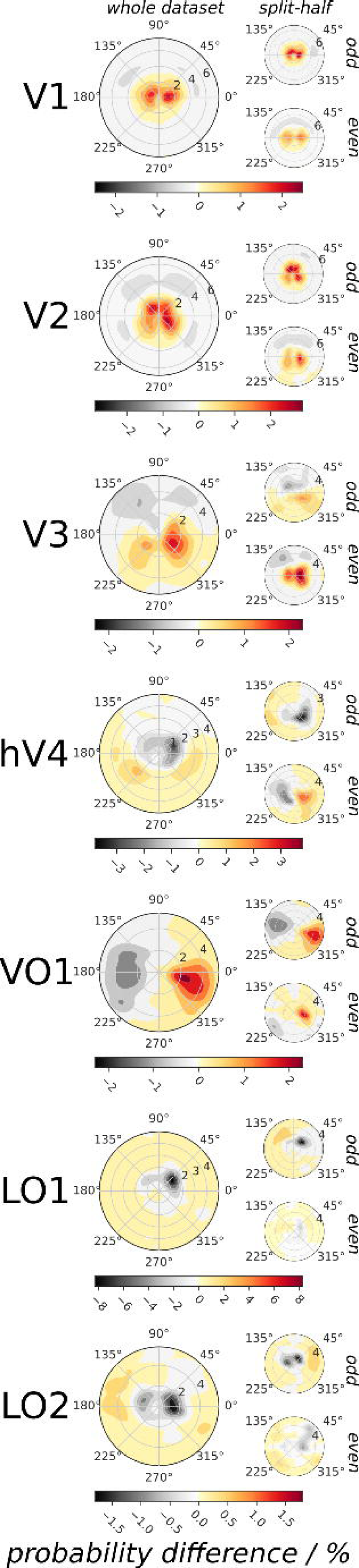
Luminance preference across the visual field. Same conventions as in Figure 10 except that preference is shown for high versus low intensity luminance. Since classification was binary, there is only a single column. Hot areas mark regions in the visual field where high luminance was preferred and gray areas mark regions where low luminance was preferred. In early areas V1-V3 for instance, luminance preference changed as a function of eccentricity. Note that although VO1 showed a difference between left and right visual field descriptively, preference classification in this area was not significant (Figure 9).

In sum, the results showed that voxels at different retinotopic locations were predicted to prefer different stimulus classes. These differences were on the order of a few percent points. These results indicate a systematic relationship between a voxel’s retinotopic spatial preference and its response pattern to color and luminance that could at least partially mediate between-subject classification.

#### ROI-specificity of SRMs for voxel preference prediction

The topographies presented in Figures 10 and 11 showed variability between the different ROIs. This raises the question to what extent the retinotopic biases in stimulus preference captured by the SRMs were unique to each visual area and to what extent they were of a more generic nature and therefore interchangeable across areas for the prediction of stimulus preference. We examined this by comparing two analyses that used SRMs trained on retinotopic data: in the first, the SRMs used to transform the training brain data into the common space and to transform the predictions into the individual space of the left-out test brain differed in terms of the ROIs they were trained on, and results from all combinations of ROIs were averaged (incongruent condition). In the second, the SRMs for both transformations were trained on the same ROI (congruent condition – identical to our original analysis).

If SRMs were specific to each visual area, one would expect that the prediction of voxel stimulus preferences is better when SRMs from the same ROI were used for both transformations (congruent condition) than when each were based on SRMs from different ROIs (incongruent condition). If the models merely capture general retinotopic biases, one would expect that prediction accuracies would not differ. As is evident from Figure 12 (left), prediction accuracies for color were significantly higher when using congruent SRM pairings than when using incongruent ones (all p < .01) in all ROIs where prediction accuracies were significantly better than chance. The only exception to this was hV4 where, although the pair-wise test was not significant (p = .1653), accuracy was significantly better than chance only for congruent pairings (p < .01) but not incongruent pairings (p = .0695). V2 was the only ROI where color preference prediction was significantly better than chance also for incongruent pairings (p < .01) while all remaining ROIs showed no significant accuracies. The same pattern of results was obtained for luminance (Figure 12, right). In all ROIs where luminance preference prediction was significantly above chance, accuracy was also significantly lower when incongruent SRM pairings were used (p < .01). Area hV4 again differed in this regard because, although the two conditions did not significantly differ (p = .2905), preference prediction accuracy was only significantly better than chance in the congruent condition (p < .01), but not the incongruent condition (p = .2218). For luminance, there was no ROI with significant prediction accuracy.

**Figure 12.**
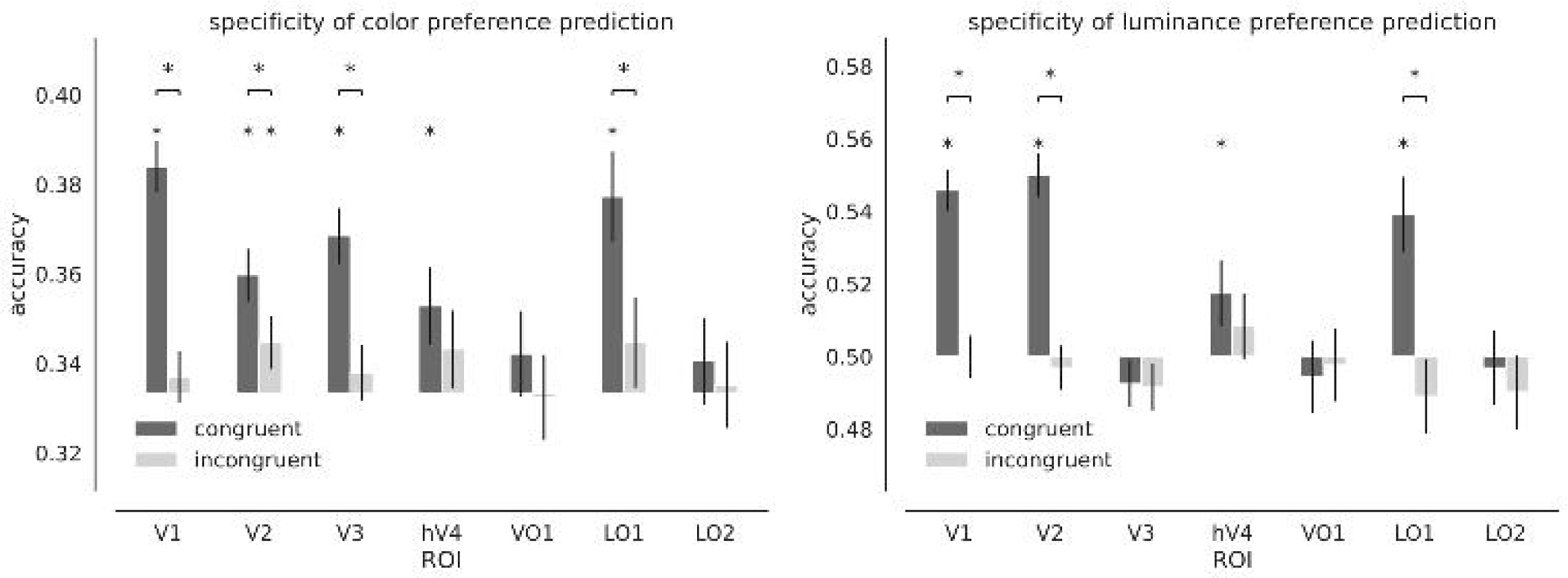
ROI-specificity of SRMs for preference prediction. To examine whether the retinotopic biases captured in SRMs were specific for individual visual areas, we repeated the voxel preference prediction analysis (Figure 8 B) but compared transformation matrices from all visual areas in the test brain with respect to how well they could predict voxel preference in a given area. Dark bars show how well voxel preference could be classified when the transformation from the SRM of the same visual area was used for prediction (“congruent”, same as mid-gray bar in Figure 9). Light bars show accuracy averaged across all classifications in which transformation into the individual space of the test brain was based on an SRM from a different visual area (“incongruent”). Accuracy is the fraction of correctly classified voxels across all brains and error bars represent 95 % CIs. Binomial tests were used to test if observed accuracies exceeded chance, z-tests were used for comparisons between congruency conditions (asterisks denote p < .05, Holm-Šidák corrected for seven ROIs).

The discrepancies in prediction accuracies between congruent and incongruent SRM pairings demonstrate that SRMs capture retinotopic biases in stimulus preferences that are largely specific to individual visual areas.

### SRM searchlight BSC analysis

We next tested how well common shared responses could be estimated from only anatomically aligned activity patterns for subsequent BSC (in contrast to our ROI BSC analysis where visual areas had been delineated functionally through retinotopy). Individual datasets were warped to MNI space separately and SRM was fit to only local patterns of retinotopic mapping responses in a searchlight analysis to estimate common space representations for every brain location. After transforming individual color responses to this common space, BSC could be carried out on those local patterns. This approach had the additional advantage that BSC could be applied to the whole brain.

Applying a false discovery rate of q < .05, we found that classification accuracies in a region within the early visual cortex was significantly larger than chance (one-tailed binomial test based on 3240 Bernoulli trials). Figure 13 shows the location of this region relative to the hV4 group ROI. This ROI comprised all the voxels that were identified as being part of individual hV4 ROIs in at least 25 % of the participants. As can be seen, the voxels where classification was significantly better than chance were located near the calcarine sulcus but did not overlap with the hV4 group ROI.

**Figure 13.**
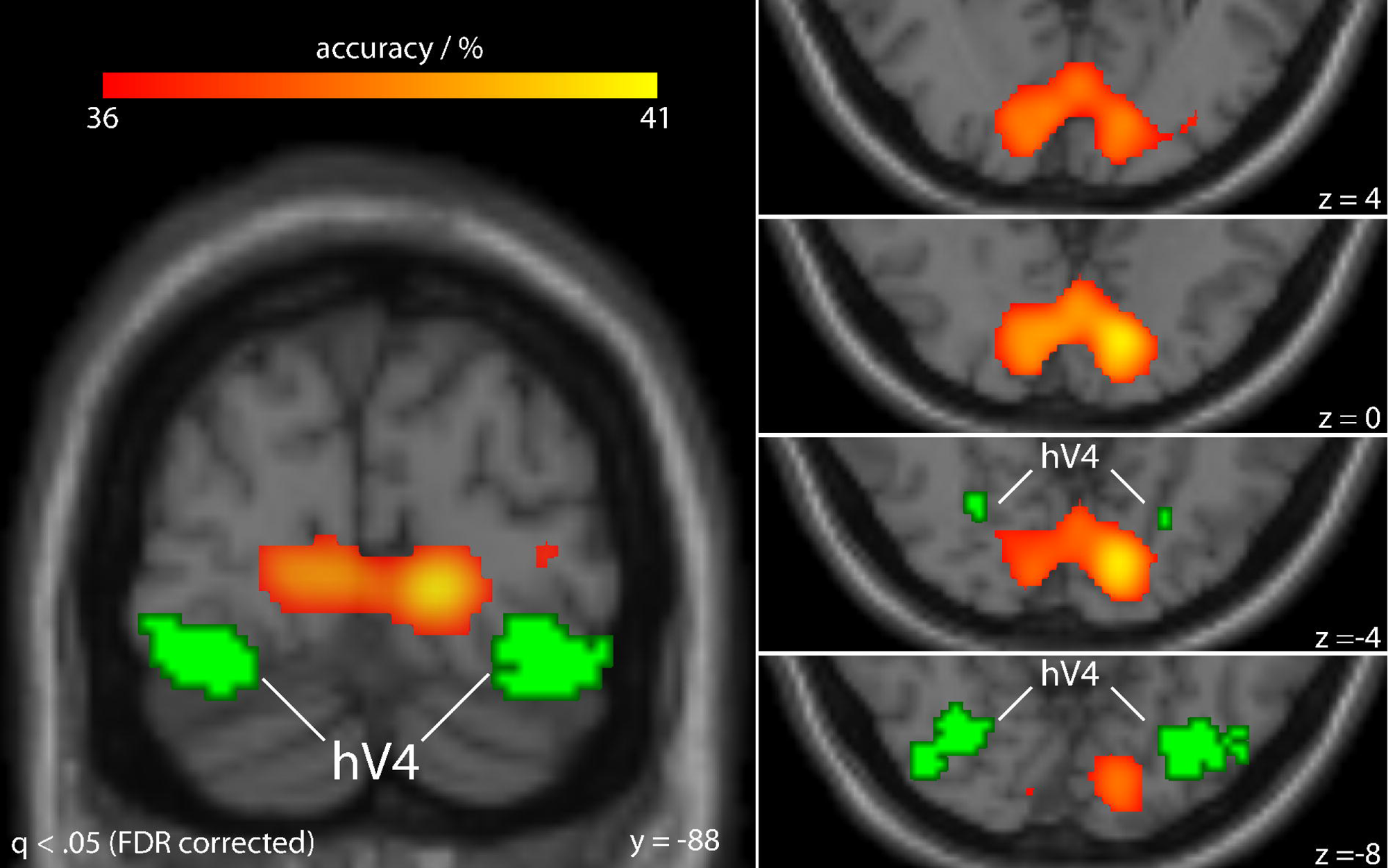
SRM searchlight map. SRMs were fit to patterns within local spheres (3 voxel radius) of brain responses to the retinotopic mapping stimulus. After transforming every participant’s dataset into the common space, classifiers were used to predict the color of the viewed image using only the information in the local activity pattern (BSC). Cross-validation was performed by leaving out data from a different participant for testing on every iteration. Classification accuracies are shown in warm colors on a standard MNI template for brain regions that survived the whole-brain significance threshold of a binomial test (q < .05, FDR corrected). Green marks all voxels falling within area hV4 in at least 25 % of the participants. The regions where BSC of color yielded accuracies significantly above chance, namely near the calcarine sulcus, were distinctly different from the location of hV4. This indicates that anatomical alignment was not sufficient beyond earliest visual areas for BSC (e.g. in hV4).

In sum, the searchlight analysis suggests that while anatomical alignment of voxels was sufficiently high for BSC of color in early visual cortex this alignment was too coarse in more anterior regions.

### Behavioral control analysis

In order to ascertain that the classification results were not driven by differences in task performance we performed a Bayesian two-way repeated-measures ANOVA (Rouder et al., 2017) of the reaction time data under the factorially crossed three-level factor color and the two-level factor luminance. Reaction time datasets from participants 2 and 9 were not complete due to programming error and had to be discarded from the analysis, yielding 13 participants in total. Residuals were normally distributed (six Shapiro-Wilk’s tests, smallest test statistic across six conditions: W = .925, p = .2931) and no violation of sphericity was detected (Mauchly’s test, p = .0845 for color, p = .6332 for the interaction). The Bayes analysis suggested that behavioral data were not affected by color or luminance: the resulting Bayes factors indicated that the data were BF_10_ = 1.273 times more compatible with the model assuming a main effect of color relative to no effect (red: M = .574 s, SD = .052 s, green M = .582 s, SD = .058 s, yellow: M = .601 s (SD = .067 s). As for luminance, the data were BF_10_ = 1.523 more likely to have occurred under the model assuming a difference between conditions compared to no difference (high: M = .595 s, SD = .054 s, low: M = .576 s, SD = .064 s). The data provided BF_01_ = 1.478 times more support to the model without interaction. Thus, even if a difference between color (and likewise luminance) conditions existed, it is too small for the present data to detect with sufficient certainty (1/3 < BF < 3 are considered “anecdotal evidence” (Jeffreys, 1961).

### Retinotopic binning analysis

Finally, we carried out a more traditional analysis that had previously been used to test whether retinotopic bias might partially underlie the accuracy with which sensory input can be decoded from patterns of fMRI signals (Beckett et al., 2012; Freeman et al., 2011; Larsson et al., 2017). We therefore repeated the WSC analysis of color yet this time binned voxels based on (a) their eccentricity preference and (b) their polar angle preference, each of which preserves to some extent the retinotopic structure in the respective coordinates. Voxel values were averaged within bins and classification was carried out on these bin averages. Accuracies were compared with accuracies obtained with random binning of voxels.

Figure 14A shows that for binning based on eccentricity, decoding accuracies were significantly better for at least some binning steps in all examined ROIs except VO1 compared to random binning. For binning based on polar angle (Figure 14B), we observed significantly higher classification accuracies in V1 and V2 only. We note that these retinotopic binning-based results are overall consistent with those of the SRM analysis shown in Figure 3. In particular, all ROIs where prediction of color from shared response patterns was significantly better than chance also showed better classification after retinotopic binning than random binning, be it based on polar angle, on eccentricity, or on both. Retinotopic binning yielded stronger results only in LO2 where classification accuracy was higher than random binning at two binning steps (after correction) whereas SRM-based results in LO2 between brains was not significantly better than chance after FWE correction across ROIs. To conclude, retinotopic binning analysis is in good agreement with the results from our BSC analysis and confirms the presence of spatial retinotopic biases in color and luminance encoding in the human brain.

**Figure 14.**
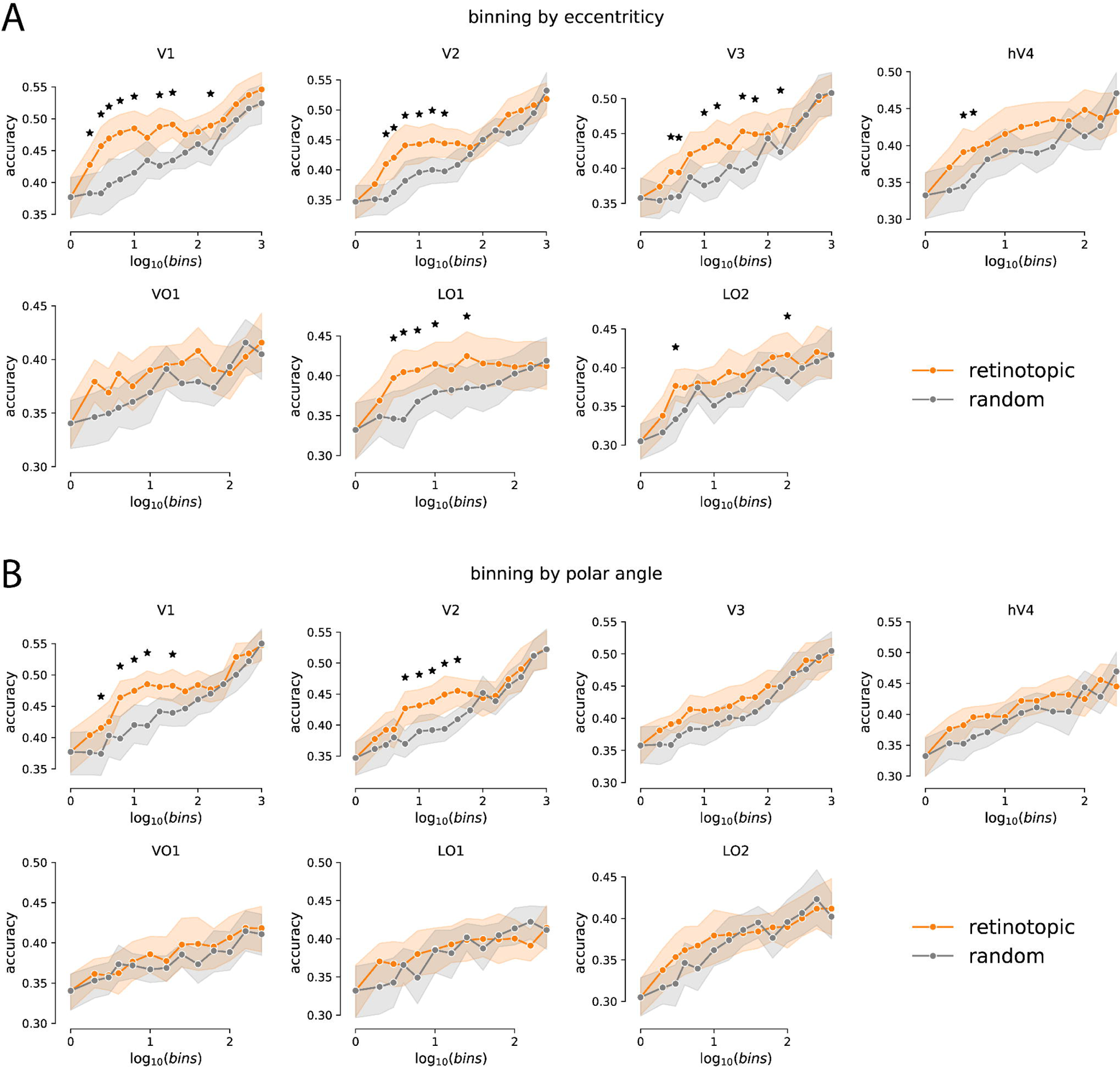
Binning analysis. Voxels within a ROI were assigned into a number of bins based on their polar coordinates, i.e., either eccentricity or polar angle. Bin limits were chosen such that they were equally spaced across the range of voxel coordinates. Different numbers of bins were used. Within-subject classification was performed except that, instead of training and testing classifiers on vectors of voxel responses, vectors of bin-wise averages of voxel responses were used. In a control analysis voxels were randomly assigned to the bins. If a retinotopic bias exists, one would expect that retinotopically based binning would yield higher classification accuracy than random binning. Paired t-tests (df = 14) were used and error bands represent 95 % bootstrapped CI. Asterisks denote significance at p < .05 (Holm-Šidák corrected for 15 binning steps). In all ROIs where color could successfully be classified across subjects in the BSC analysis also the binning analysis also showed a retinotopic bias when binning based on eccentricity or polar angle (or both, see V1 and V2), thus confirming the results from our own analysis. In addition, retinotopic binning yielded higher accuracies than random binning also in LO2.

## Discussion

Shared response modeling allows experimenters to test for between-subject commonalities in the neural representations of sensory stimuli. Typical applications involve sampling large neural response spaces from individual observers by presenting them with a vast variety of stimulus input (e.g., movies). Knowing how different brains respond to such stimulation makes it possible to match idiosyncratic response spaces to each other and to identify neural representations that are shared across observers. We applied SRM to brain activity measured while observers viewed simple retinotopic mapping stimuli, not movies. We show that controlling for how different brains respond to the same, purely spatially defined mapping stimulus is sufficient to preserve color responses across brains while being specific for distinct areas across the visual processing hierarchy. Our results suggest a relationship between visual space and color that is shared across observers and unique to each visual area. This conclusion is supported by three key observations.

First, we fit SRMs to retinotopic mapping data to map individual fMRI patterns into a shared common space. Classification analysis in this space showed that, in areas V1, V2, V3, hV4, and LO1, it was possible to predict what color an observer was seeing even though classifiers were trained exclusively on responses from other observers’ brains (Figure 3). We found no behavioral differences between stimulus conditions that could explain these results. When constraining our analysis to subsets of voxels, V1, hV4, and LO1 showed the largest degree of robustness across voxel selections in accordance with their important role for color vision (Bartels & Zeki, 2000; Beauchamp et al., 1999, p. 1999; Brouwer & Heeger, 2009; Engel et al., 1997; Lueck et al., 1989). In conclusion, this demonstrates that aligning individual response spaces across brains with respect to their retinotopic functional organization is sufficient to afford classification of color in multiple visual areas across different brains.

Second, we hypothesized that the systematic relationship between retinotopic and color responses captured by SRMs would allow them to predict how individual voxels would respond to the color stimuli in an fMRI dataset recorded from a completely independent brain. Similar methods had previously been used to predict broader brain topographies like the location of FFA and PPA (Haxby et al., 2011). SRM-based prediction accuracies at this task were indeed above chance in various ROIs (Figure 9) although no consistent preference pattern is discernible when inspecting color preferences of voxels projected onto individual inflated hemispheres (Figures 4 and 5). Similarly, a previous study presenting “color-preference images (similar to the orientation-preference maps of Kamitani and Tong (2005)” found no evidence for regional bias either (Figure 11, Parkes et al., 2009). This seemingly contradicts our observation that voxel preferences could be predicted across brains. This disparity is easily explained if one considers that SRM separates individual participants’ fMRI responses into shared and idiosyncratic components. While retinotopic biases exist in the shared component, it is conceivable that at the individual level they remain obscured by idiosyncratic response characteristics. Likewise, retinotopic biases as an explanation of orientation decoding (Freeman et al., 2011) were demonstrated with an analysis like our binning test (Figure 14) whereas they had not been apparent from individual weight maps (Kamitani & Tong, 2005).

Third, we quantitatively investigated if the relationships between retinotopy and color responses captured by SRMs were specific to different ROIs. We therefore compared how accurately voxel responses to color could be predicted when transformation matrices from SRMs belonging to either the same or different areas were used for model fitting and prediction, respectively. We found that prediction accuracies dropped considerably when SRMs based on different visual areas were used (Figure 12). This specificity is consistent with visual inspection showing marked differences between topographies from different brain areas (Figure 10). These inter-area differences are important as they rule out alternative explanations like artefactual spatial inhomogeneities, e.g., in the stimulus projection, because, in that case, one would expect identical topographies across ROIs.

The decreasing preference for red and green stimuli in V1 at larger eccentricity is consistent with previous fMRI findings reporting decreasing responsiveness to modulations in the red/green direction in color space, but not for the blue/yellow direction (Mullen et al., 2007; Vanni et al., 2006). Similarly, behavioral measurements showed that, compared to blue-yellow and achromatic mechanisms, contrast sensitivities of the red-green cone mechanism are highest at the fovea, yet show the strongest decline as eccentricity increases (Hansen et al., 2009; Mullen, 1991; Mullen & Kingdom, 2002). Although our stimuli did not systematically sample color space and did not include blue, our V1 findings are compatible with behavior. Such eccentricity differences may be related to the observation that cone density decreases with eccentricity (Curcio et al., 1990a), or that S cones are not found in the fovea (Roorda & Williams, 1999), or that a fourth (melanopsin) photopigment may contribute to peripheral color vision (Horiguchi et al., 2013). However, the distributions of L and M cones (in contrast to S cones) show large variability across individuals (Brainard, 2015) so that commonalities at the receptor level likely cannot fully explain the cortical effects.

V2, unlike V1, shows no preference for green in the fovea but only to the left and the right of it around the horizontal meridian while green is relatively less preferred along the upper vertical meridian and yellow is less preferred along the horizontal meridian. In V3, the fovea has less preference for green while, in every quadrant, there is a patch each surrounding it where green is preferred. The opposite pattern is found for red and, to lesser extent, for yellow, which are both more preferred in the center and more peripheral regions of the visual field. Area hV4 shows higher preference for red than green or yellow in the center, which drops off ventrally and to its left and right where yellow and green are preferred. Finally, LO1 shows more heterogeneity in preference in the upper visual field. These descriptive differences between visual areas are consistent with our finding that stimulus preference can be predicted more accurately when SRMs from the same visual area are used to train models and derive predictions.

Similar variability between areas has been observed using fMRI with respect to spatial sensitivity for chromatic and achromatic input which differentially covaries with eccentricity (Welbourne et al., 2018). In addition to eccentricity, human neuroimaging studies – although not specifically addressing color vision – demonstrated that polar angle influences visual processing (Benson et al., 2012; Liu et al., 2006; Silson et al., 2018; Silva et al., 2018). It is hence conceivable that the topographies in extrastriate areas are related to different behavioral and cone mechanisms than V1, involving a combination of both eccentricity and polar angle differences. While it is known that visual perception of spatial frequency and orientation depends on polar angle (Jóhannesson et al., 2018; Karim & Kojima, 2010), the empirical findings for color are mixed (Danilova & Mollon, 2009; Levine & McAnany, 2005). At the retinal level, cone densities change depending on polar angle in that they are higher along the horizontal than the vertical meridians (Curcio et al., 1990b; Song et al., 2011). But at least for orientation discrimination, simulations suggested that these cone differences cannot fully explain that polar angle affects visual performance (Kupers et al., 2019).

The color biases presented here complement similar fMRI findings involving other visual features like orientation (Beckett et al., 2012; Freeman et al., 2011, 2013; Larsson et al., 2017; Sasaki et al., 2006) and we predict that retinotopy-based SRM would also allow between-subject classification of these features. Concerning color, spatial biases in the visual system are known from research on mice in which stimuli appearing above or below the horizon carry different behavioral meaning (Baden et al., 2013; Denman et al., 2018; Rhim et al., 2017). Environmental regularities like the chromaticities of natural daylight (Lafer-Sousa et al., 2012) or of objects versus their backgrounds (Rosenthal et al., 2018) shape color vision in primates, too. Which environmental adaptations may underlie our findings requires further research.

We point out that our stimuli do not only probe the retinotopic layout of cortical field maps. That is because they are composed of more low-level spatial features like contrast, motion, and orientation, which may have shared representations with color (Friedman et al., 2003; Garg et al., 2019; Johnson et al., 2001, 2008; Leventhal et al., 1995; Sincich & Horton, 2005). SRM may therefore also exploit commonalities in the co-representations of these features, thereby contributing to our results.

Finally, it has been demonstrated that response biases for orientation can be changed by embedding a grating stimulus in a modulator, which could render the orientation either radial or tangential (Roth et al., 2018). Color vision, on the other hand, shows some robustness in that different colors elicit patterns of brain activity that are similar between sensory-driven colors and memory colors (Bannert & Bartels, 2013; Teichmann et al., 2019; Vandenbroucke et al., 2016) and across tasks (Bannert & Bartels, 2018; Serences et al., 2009). In sum, given the evidence for both flexibility and stability of color representations, it would be interesting for future studies to probe the plasticity of the observed large-scale response biases.

## Acknowledgments

This work was supported by the German Excellence Initiative of the German Research Foundation (DFG) grant number EXC307, by the Max Planck Society, Germany, and by DFG grant SFB 1233, Robust Vision: Inference Principles and Neural Mechanisms, project 9, project number: 276693517.

